# The T1150A cancer mutant of the protein lysine dimethyltransferase NSD2 can introduce H3K36 trimethylation

**DOI:** 10.1101/2023.03.13.532367

**Authors:** Mina S. Khella, Philipp Schnee, Sara Weirich, Tan Bui, Alexander Bröhm, Pavel Bashtrykov, Jürgen Pleiss, Albert Jeltsch

## Abstract

Somatic mutations in protein lysine methyltransferases are frequently observed in cancer cells. We show here that the NSD1 mutations Y1971C, R2017Q and R2017L observed mostly in solid cancers are catalytically inactive suggesting that NSD1 acts as tumor suppressor gene in these tumors. In contrast, the frequent T1150A in NSD2 and its T2029A counterpart in NSD1, both observed in leukemia, are hyperactive and introduce up to H3K36me3 in biochemical and cellular assays, while wildtype NSD2 and NSD1 only generate up to H3K36me2. MD simulations with NSD2 revealed that H3K36me3 formation is possible due to an enlarged active site pocket of T1150A and loss of direct contacts of T1150 to critical residues which regulate the product specificity of NSD2. Bioinformatic analyses of published data suggest that the NSD2 T1150A mutation in lymphocytic leukemia could alter gene regulation by antagonizing H3K27me3 finally leading to the upregulation of oncogenes.

## 1 Introduction

Epigenetic modifications including DNA methylation and histone modifications are key determinants of important cellular processes and differentiation events (Allis and Jenuwein, 2016; Jambhekar et al., 2019) and perturbations in epigenetic mechanisms can lead to initiation and progression of many diseases including cancer (Zhao et al., 2021). Many different types of histone post-translational modifications have been discovered so far, including lysine and arginine methylation, lysine acetylation, threonine and serine phosphorylation and others (Huang et al., 2015). The combination of all these histone modifications, forming the so-called histone code, directs the organization of chromatin and subsequent control of gene expression. Lysine methylation of histone N-terminal tails is one of the most abundant histone modifications which is introduced by the site-specific protein lysine methyltransferases (PKMTs) (Husmann and Gozani, 2019). Depending on the site of the methylated lysine, it could be linked either with repressive (H3K9me2/3 and H3K27me3) or active chromatin states (as in case of H3K4me1/2/3). Histone H3 lysine 36 (H3K36) methylation is connected with several biological processes including control of gene expression, DNA repair and recombination, as well as alternative splicing (Lam et al., 2022; Li et al., 2019; Wagner and Carpenter, 2012). H3K36 methylation in human cells is mainly catalyzed by 5 different enzymes (NSD1, NSD2, NSD3, ASH1L and SETD2). While NSD1, NSD2, NSD3 and ASH1L can introduce only up to dimethylation of H3K36 in vitro and in vivo (Husmann and Gozani, 2019; Li et al., 2019; Li et al., 2009; Qiao et al., 2011), SETD2 is the main human enzyme so far which can introduce up to trimethylation of H3K36 (H3K36me3) in somatic cells (Edmunds et al., 2008; Eram et al., 2015). The difference in the methylation states together with their different genomic localization are key factors in the complexity of the H3K36 histone code, because each methylation state of K36 encodes different biological downstream cascades (DiFiore et al., 2020; Li et al., 2019; Wagner and Carpenter, 2012). The H3K36me2 mark occurs at intergenic regions and promotors while H3K36me3 mark is enriched at gene bodies of active genes (Cornett et al., 2019; Hyun et al., 2017; Yun et al., 2011; Zhou et al., 2011). H3K36me2 is directly connected with the DNA methyltransferase DNMT3A via its the proline-tryptophan-tryptophan-proline (PWWP) domain of which binds preferentially to H3K36me2 at intergenic regions (Dhayalan et al., 2010; Dukatz et al., 2019; Weinberg et al., 2019), while H3K36me3 is bound by the PWWP domain of DNMT3B (Baubec et al., 2015). H3K36me2/3 were shown to act as antagonists of H3K27 trimethylation (Finogenova et al., 2020; Streubel et al., 2018), but this effect is more pronounced in case of H3K36me3, as illustrated by global mass spectrometric analyses (Leroy et al., 2013; Mao et al., 2015; Voigt et al., 2012). H3K27me3 is another repressive histone mark which is deposited by PRC2 complex and has important role in development.

The nuclear receptor binding SET domain protein 2 (NSD2) (also known as MMSET, and WHSC1) (Lam et al., 2022) and its human paralogs NSD1 (also known as KMT3B) (Tauchmann and Schwaller, 2021) and NSD3 (WHSC1L1) (Rathert, 2021), are Su(var)3–9, Enhancer-of-zeste, Trithorax (SET) domain containing PKMTs (Lam et al., 2022; Rayasam et al., 2003). All NSD enzymes share a similar domain organization. Their catalytic SET domain at the C-terminal part of the enzyme comprises 3 subdomains, pre-SET or associated with SET (AWS), SET, and post-SET. It is responsible for methylation of histone H3K36 using the methyl donor cofactor S-adenosyl-L-methionine (AdoMet) (Lam et al., 2022; Rathert, 2021). Similar to some other PKMTs (for example Clr4 and EZH2 (Iglesias et al., 2018; Khella et al., 2020; Lee et al., 2019; Wang et al., 2019)), NSD PKMTs adopt an autoinhibitory conformation in which an autoregulatory loop (ARL) connecting the SET and post-SET domains blocks the substrate lysine binding channel in the absence of substrate (Qiao et al., 2011; Tisi et al., 2016). Recently, a crystal structure of NSD2 bound to a nucleosomal substrate revealed crucial contacts of NSD2 with the DNA, the H3 tail and histone H2A which altogether stabilize the NSD2 ARL in an active open conformation (Li et al., 2021; Sato et al., 2021).

In addition to the catalytic SET domain, NSD enzymes contain other regulatory domains including two PWWP domains which are important for binding to DNA and methylated H3K36. In case of NSD2, the first N-terminal PWWP domain was shown to bind H3K36me2 and stabilizes NSD2 binding at chromatin (Sankaran et al., 2016). Furthermore, the NSD enzymes contain five plant homeodomains (PHD) and an atypical (C5HCH) plant homeodomain (PHD) finger and NSD2 has a unique high mobility group (HMG) motif which contributes to its nuclear localization (Kang et al., 2009; Lam et al., 2022). In the context of nucleosomal substrates, H3K36 was the specific site to be methylated by NSD2 (Li et al., 2009). In case of NSD1, K168 of the linker histone H1.5 was shown to be methylated in addition to H3K36 (Kudithipudi et al., 2014b). Moreover, some non-histone targets were shown to be methylated by NSD1, like NFkB-p65 (K218 and K221) (Lu et al., 2010), ATRX (K1033) as well as the small nuclear RNA-binding protein U3 (K189) (Kudithipudi et al., 2014b).

In agreement with the complex functions of NSD-catalyzed H3K36 dimethylation and non-histone methylation, NSD enzyme dysfunction was linked to several diseases ranging from developmental disorders to cancers (Lam et al., 2022; Li et al., 2019; Wagner and Carpenter, 2012). Heterozygous loss of NSD2 is responsible for a developmental disease called Wolf-Hirschhorn syndrome (WHS) (Bergemann et al., 2005). Haploinsufficieny of NSD1 was linked to an overgrowth syndrome called SOTOS syndrome (Kurotaki et al., 2002). Regarding cancer involvement, NSD2 is overexpressed and acts as major regulator of gene transcription and disease progression in multiple myeloma cells harboring t (4;14) translocation (Martinez-Garcia et al., 2011). Increased activity of NSD1 is a common feature in AML with the t(5;11)(q35;115) chromosomal translocation which results in a fusion of the N-terminal domains of the nucleopore 98 (NUP98) protein to the C-terminal part of NSD1 (Hollink et al., 2011; Jaju et al., 2001; Wang et al., 2007). In addition, many missense variants of NSD1/2 were observed in various types of cancers like hematological cancers (Jaffe et al., 2013; Leonards et al., 2020), head and neck squamous cell carcinomas (HNSCC) (Network, 2015), human brain tumor cell lines (Berdasco et al., 2009), and lung cancers (Brennan et al., 2017; Sengupta et al., 2021; Yuan et al., 2021).

Unlike the loss-of-function changes caused by gene deletions, deciphering the biological effects caused by somatic cancer missense mutation is more challenging. Many mutations of this type are observed in PKMTs in different types of cancers and some of them were shown to dramatically change enzyme activity, product specificity, substrate specificity or other enzyme properties (Bröhm et al., 2019; Oyer et al., 2014; Weirich et al., 2017; Weirich et al., 2015). This highlights how a single amino acid exchange could mechanistically drive carcinogenesis. A frequent NSD2 missense single point mutation (E1099K) was detected in many leukemic patients. This mutant was comprehensively characterized and shown to be hyperactive (Jaffe et al., 2013; Oyer et al., 2014; Pierro et al., 2020; Swaroop et al., 2019). Structural analyses revealed that it destabilizes the NSD2 ARL leading to higher activity (Li et al., 2021; Sato et al., 2021). However, the effects of other frequent missense cancer mutants in NSD2 and its paralog NSD1 are not understood. In particular, T1150A NSD2 is one of the frequent somatic cancer mutants and the corresponding T1232A mutation in NSD3 has been shown before to be a driver mutant which increases cell proliferation and xenograft tumor growth (Li et al., 2021)

In this work, we aimed to characterize the biochemical effects of the frequent somatic missense cancer mutation in NSD2 (T1150A) and its NSD1 analogue (T2029A) both observed in leukemia in addition to three other somatic cancer mutations in NSD1 observed mostly in solid cancers (Y1971C, R2017Q and R2017L). We show that Y1971 and R2017 are critical residues in NSD1 and their mutation leads to a loss of enzyme activity suggesting that NSD1 acts as tumor suppressor gene in these cancers. In contrast, our data reveal that the T1150A/T2029A mutants of NSD2 and NSD1 are hyperactive and in biochemical and cellular assays, we observed that they introduce up to H3K36me3, while the wildtype (WT) NSD2 and NSD1 enzymes can only generate H3K36me2. In Molecular Dynamics (MD) simulations we demonstrate that H3K36me3 generation is possible due to an enlarged active site pocket of the mutant that allows AdoMet binding to an NSD2 T1150A-H3K36me2 complex in a productive conformation leading to H3K36me3 generation. Furthermore, we provide evidence that the generation of H3K36me3 leads to altered gene regulation in lymphocytic leukemia tumor cells by antagonizing H3K27me3 finally leading to the upregulation of oncogenes.

## 2 Materials and Methods

### 2.1 Site-directed mutagenesis, enzymes overexpression and purification

The GST-tagged expression constructs of human NSD2 catalytic domain including the AWS, SET and Post-SET domains (amino acids 992–1240 of UniProt No: O96028) was prepared. This part is an independent folding unit as illustrated by structural studies [pdb 7E8D (Sato et al., 2021)]. A corresponding GST-tagged mouse NSD1 catalytic domain (amino acids 1701–1987, UniProt No: O88491) containing an additional C1920S mutation was taken from (Kudithipudi et al., 2014a). The SET domains of human and mouse NSD1 share >95% identity and none of the residues that are affected in this study or known to be catalytically relevant resides is different. The different NSD2 (T1150A) and NSD1 mutants (Y1971C, R2017Q/L and T2029A) were created by site-directed mutagenesis using the megaprimer method (Jeltsch and Lanio, 2002). Human NSD1 residues Y1971, R2017 and T2029 correspond to mouse NSD1 residues Y1869, R1915 and T1927. The sequence of all plasmids was validated by Sanger sequencing. For the protein overexpression, the different plasmid constructs (WT and mutants) were transformed into *E. coli* BL21-CodonPlus (DE3) cells (Novagen). The bacterial cells were grown at 37 °C until they reached an OD^600nm^ between 0.6 and 0.8. Afterwards, 1 mM isopropyl-β-d-thiogalactopyranoside (IPTG) was added to induce protein expression at 20 °C overnight. The next day, the cells were harvested by centrifugation at 3800 rcf for 20 min, followed by washing once with STE buffer (10 mM Tris/HCl pH 8.0, 1 mM EDTA and 100 mM NaCl) and collection of the cell pellets by centrifugation at 4900 rcf for 25 min.

For NSD1 and NSD2 protein purification, GST-tag affinity chromatography was used. In brief, the cell pellets were thawed on ice and resuspended in sonication buffer (50 mM Tris/HCl pH 7.5, 150 mM NaCl, 1 mM DTT, 5% glycerol) supplemented with protease inhibitor cocktail containing AEBSF-HCL (1 mM, Biosynth), pepstatin (10 μM, Roth), aprotinin (0.4 μM, Applichem), E-64 (15.14 μM, Applichem), leupeptin (22.3 μM Alfa Aesar) and bestatin (50 μM, Alfa Aesar) and the cells were disrupted by sonication. The lysed cells were centrifuged at 40,000 rcf for 90 min at 4 °C. The supernatant was loaded onto a column containing sonication buffer pre-equilibrated glutathione-Sepharose 4B beads resin (GE Healthcare). Afterwards, the beads were washed once with sonication buffer and twice with wash buffer (50 mM Tris/HCl pH 8, 500 mM NaCl, 1 mM DTT, 5 % glycerol) and the bound proteins were eluted with the wash buffer containing 40 mM reduced Glutathione. Fractions containing the protein were pooled and proceeded to dialysis against first dialysis buffer (20 mM Tris/HCl pH 7.2, 100 mM KCl, 0.5 mM DTT, 10% glycerol) for 3 hours followed by second dialysis against a second dialysis buffer (20 mM Tris/HCl pH 7.2, 100 mM KCl, 0.5 mM DTT, 60% glycerol) overnight. The protein solution was stored in aliquots at −20 °C. Purified proteins were analyzed by sodium dodecyl sulfate–polyacrylamide gel electrophoresis (SDS-PAGE) using 16% gels stained with colloidal Coomassie brilliant blue.

Histone octamers were prepared as described previously (Bröhm et al., 2022). Briefly, the pET21a expression constructs of H3.1, H4, H2A and H2B were overexpressed in BL21-Codon Plus *E. coli* cells which were allowed to grow at 37 °C until OD^600nm^ of 0.6–0.8 was reached. Induction of protein expression was done at 20 °C shaker for 3 h after the addition of. Cells were harvested by centrifugation at 5000 rcf for 15 min, washed using STE buffer (10 mM Tris/HCl pH 8, 100 mM NaCl, 1 mM EDTA), centrifuged again at 5000 rcf for 15 min and the cell pellets stored at −20 °C.

For histone protein purification, the bacterial cell pellets were resuspended in SAU buffer (10 mM sodium acetate pH 7.5, 1 mM EDTA, 10 mM lysine, 5 mM β-mercapthoethanol, 6 M urea, 200 mM NaCl) followed by sonication (Epishear, Active Motif). The lysate was centrifuged at 40,000 rcf for 1 h and the supernatant was filtered through a 0.45 μM syringe filter (Chromafil GF/PET 45, MachereyNagel) and passed through a HiTrap SP HP (5 mL, GE Healthcare) column connected to an NGC FPLC system (BioRad), which was previously equilibrated with SAU buffer. After washing the column with SAU buffer, proteins were eluted with NaCl gradient from 200 mM to 800 mM. Fractions were collected, analysed by SDS-PAGE, pooled according to purity and yield, dialyzed against pure water with two changes overnight and dried in a vacuum centrifuge for storage at 4 °C.

The lyophilized histone proteins were dissolved in unfolding buffer (20 mM Tris/HCl pH 7.5, 7 M guanidinium chloride, 5 mM DTT) and their concentrations were determined spectrophotometrically at OD^280nm^. The proteins were mixed in a ratio of 1 (H3, H4) to 1.2 (H2A, H2B). The samples were dialyzed against refolding buffer (10 mM Tris/HCl pH 7.5, 1 mM EDTA, 2 M NaCl, 5 mM β-mercapthoethanol) overnight with one buffer change. To purify the octamers afterwards, the samples were separated by size exclusion chromatography using a Superdex 200 16/60 PG column equilibrated with refolding buffer. Fractions were collected, pooled according to purity and afterwards concentrated using Amicon Ultra-4 centrifuge filters (30 kDa cutoff, Merck Millipore). The purified octamers were validated by SDS-PAGE, aliquoted, flash frozen in liquid N2 and stored at −80 °C.

### 2.2 Nucleosome reconstitution

Nucleosomes were prepared using the histone octamers and DNA fragments as described previously (Bröhm et al., 2022). Briefly, the Widom-601 sequence (Lowary and Widom, 1998) was cloned into a TOPO-TA vector together with a linker sequence providing 64 bp linker DNA on the 5’ side of the core nucleosome and 29 bp on the 3’ side, amplified by PCR and purified. DNA and histone octamers were mixed in different ratios between equimolar and 2-fold octamer excess. The samples were then dialyzed in a Slide-A-Lyzer microdialysis devices (ThermoFisher) against high salt buffer (10 mM Tris/HCl pH 7.5, 2 M NaCl, 1 mM EDTA, 1 mM DTT), which was continuously replaced by low salt buffer (same as high salt, but with 250 mM NaCl) over 24 h. Afterwards, the samples were dialyzed overnight against storage buffer (10 mM Tris/HCl pH 7.5, 1 mM EDTA, 1 mM DTT, 20% glycerol), aliquoted, flash frozen in liquid nitrogen and stored at −80 °C. The DNA assembly with histone octamer in the nucleosome was validated by the EMSA (electro-mobility gel shift assay).

### 2.3 Circular Dichroism spectroscopy

The Circular Dichroism (CD) spectroscopy measurement was performed to investigate the secondary structure composition of the NSD proteins. Protein samples were mixed with 200 mM KCl buffer and CD spectra were recorded from 240 nm to 190 nm at 20 °C for 60 cycles using a 0.1 mm cuvette (J-815-150S CD Spectrometer, Jasco). As background signal, dialysis buffer II (20 mM Tris/HCl pH 7.2, 100 mM KCl, 0.5 mM DTT, 60% glycerol) was measured under the same conditions.

### 2.4 Peptide methylation assay using radioactively labelled AdoMet

NSD1/2 WT and different cancer mutants (3.4 μM) were mixed with the unmodified H3 (26–44) peptide (4.4 μM) or H1.5 (160-176) peptide (9.8 μM) (Intavis AG, Köln, Germany) in methylation buffer (50 mM Tris/HCl HCl, pH 9, 5 mM MgCl2 and 1 mM DTT) supplemented with 0.76 μM radioactively labelled AdoMet (PerkinElmer) for 4 h at 37 °C or overnight at 25 °C. The reactions were stopped by the addition of SDS-PAGE loading buffer and heated for 5 min at 95 °C. Afterwards, the samples were separated by Tricine-SDS-PAGE followed by the incubation of the gel in amplify NAMP100V (GE Healthcare) for 1 h on a shaker and drying of the gel for 2 h at 70 °C under vacuum. The signals of the transferred radioactively labelled methyl groups were detected by autoradiography using a Hyperfilm™ high performance autoradiography film (GE Healthcare) at −80 °C in the dark. The film was developed with an Optimax Typ TR machine after different exposure times. Quantification of images was conducted with ImageJ.

### 2.5 Protein and nucleosome methylation assay using radioactively labelled AdoMet

NSD1/2 WT and different cancer mutants (3.4 μM) were mixed with recombinant H3.1 protein (1 μg) (purchased from NEB) or recombinant H3.1 mononucleosomes in methylation buffer (50 mM Tris/HCl, pH 9, 5 mM MgCl2 and 1 mM DTT) supplemented with 0.76 μM radioactively labelled AdoMet (PerkinElmer) for 4 h at 37 °C or overnight at 25 °C. The reactions were stopped by the addition of SDS-PAGE loading buffer and heated for 5 min at 95 °C. Afterwards, the samples were resolved by 16% SDS-PAGE and processed as described in the last chapter for Tricine-SDS-gels.

### 2.6 Analysis of peptide methylation by MALDI mass spectrometry

The methylation reactions were performed using the unmodified H3K36 or H3K36me1 (26–44) peptides and H1.5 (160-176) (4.5 μM) in methylation buffer (50 mM Tris/HCl, pH 9, 5 mM MgCl2 and 1 mM DTT) supplemented with 1 mM unlabeled AdoMet (Sigma-Aldrich) and 6.7 μM NSD1 for 4 h at 37 °C. The reactions were halted by the addition of 0.1% trifluoroacetic acid (TFA). All the samples were cleaned using C18 tips (Agilent Technologies). The eluted samples were spotted onto an anchor chip plate (Bruker-Daltonics) followed by drying. Next, 1 μL of HCCA matrix (0.7 mg/mL α-cyano-4 hydroxycinnamic acid dissolved in 85% acetonitrile, 0.1% TFA, 1 mM ammonium dihydrogen phosphate) was added to the dried sample spots and allowed to dry again. Afterwards, the dried spots on the anchor plate were analyzed using an Autoflex Speed MALDI-TOF mass spectrometer (Bruker-Daltonics). The mass spectra were collected using the Flex control software (Bruker-Daltonics). For calibration, the peptide calibration standard (Bruker-Daltonics) with peptides ranging from 700 to 3200 Da was used. The collected spectra were then analyzed with Flex analysis software (Bruker-Daltonics).

### 2.7 Detection of H3.1 methylation by western blot

NSD1 and NSD2 WT and their corresponding cancer mutants (3.4 μM) were mixed with either recombinant H3.1 protein or recombinant H3.1 mononucleosomes in methylation buffer (50 mM Tris/HCl, pH 9, 5 mM MgCl2 and 1 mM DTT) supplemented with 1 mM unlabeled AdoMet (Sigma-Aldrich) overnight (14 h) at 25 °C. The reactions were stopped on the next day by the addition of SDS-PAGE loading buffer and heating for 5 min to 95 °C. Afterwards, the samples were resolved by 16% SDS-PAGE. Analysis was done by western blotting using the primary antibodies directed against H3K36me3 (ab9050, 1:2000) or H3K36me2 (ab9049, 1:2500) and as secondary antibody the anti-rabbit HRP (Na934v, GE Healthcare, 1:5000). The signal was detected by chemiluminescence after the addition of Pierce™ ECL Western Blotting substrate.

The antibodies used in the Western-Blot analysis were validated by peptide array binding. Peptide arrays were synthesized with an Autospot peptide array synthesizer (Intavis AG, Köln, Germany) using the SPOT method (Kudithipudi et al., 2014a; Weirich and Jeltsch, 2022). Four spots corresponding to 4 different K36 methylation states of the H3K36 peptide (29-43) (unmodified-H3K36me1-H3K36me2-H3K36me3) and one additional spot with H3K36A as negative control were synthesized on the array. After blocking with 5% milk in TBST buffer, the array was incubated with the primary antibody solution for 1 h at room temperature followed by washing 3 times. Then, the array was kept in a secondary antibody solution (anti-rabbit HRP) for 1 h. After washing again, the signal was detected by chemiluminescence after the addition of Pierce™ ECL Western Blotting substrate.

### 2.8 Cell cultivation

Authenticated HEK293 cells (RRID: CVCL_0045) were obtained from DSMZ (https://www.dsmz.de/). HEK293 cells were grown in Dulbecco’s Modified Eagle’s Medium (Sigma) supplemented with 10% fetal bovine serum, penicillin/streptomycin, and L-glutamine (Sigma) and maintained at 37 °C with 5% CO_2_.To exclude mycoplasma contamination, the cells were routinely stained with DAPI and absence of mycoplasma DNA was shown by qPCR. All experiments were conducted in mycoplasma free cell lines.

### 2.9 Preparation of SETD2 knockout HEK293 cells

As a first step, a CRISPR-Cas9 knock out of SETD2 was conducted in HEK293 cells and single cells were isolated by cell sorting. Three gRNAs were used to target the *Setd2* gene aiming to increase the probability of successful knockout as described (Chen et al., 2017) (Supplementary Table 1). Single stranded oligonucleotides encoding these sequences as forward strand were annealed to their complementary oligonucleotides to result in double stranded DNA with 5’ single stranded overhangs complementary to the BbsI restricted Cas9 vector. After that, these double stranded oligonucleotides were ligated with pU6-(BbsI)_CBh-Cas9-T2A-mCherry plasmid backbone (provided by Ralf Kuehn, Addgene catalog number 64324) (Chu et al., 2015) in presence of BbsI HF restriction enzyme (NEB) and T4 DNA ligase (NEB) using golden gate assembly. The ligated products were transformed into XL1 blue bacterial cells by electroporation followed by isolating the plasmids from single bacterial colonies using NucleoBond Xtra Midi kit (Macherey-Nagel). The Cas9-sgRNA plasmids were validated by BbsI restriction analysis and confirmed by Sanger sequencing. Next, a mixture of the 3 Cas9-sgRNA plasmids (150ng/μl) was transfected into HEK293 cells at 70% confluency in fresh medium using FuGENE^®^ HD Transfection Reagent (Promega). Two days after transfection, mCherry positive HEK293 cells were sorted using Sony cell sorter SH800S into 96 well plates containing Dulbecco’s Modified Eagle’s Medium (Sigma) supplemented with 20% fetal bovine serum, penicillin/streptomycin, and L-glutamine (Sigma). The single cell clones were allowed to grow and expand and then selected for SETD2 knockout using both H3K36me3 western blot and Sanger sequencing. Since SETD2 is the sole human enzyme responsible for depositing H3K36me3, the genomic H3K36me3 levels from the SETD2 knockout HEK293 cells were tested with H3K36me3 specific antibody (ab9050) and compared to the parental cells. For Sanger sequencing, genomic DNA was extracted from the parental and knockout HEK293 cells using QIAamp DNA mini kit (QIAGEN) and PCR amplified with primers located ~500-600 bp from the sgRNA target site. PCR products were Sanger sequenced, and sequences were aligned using the SnapGene multiple sequence alignment tool.

### 2.10 Transfection of HEK293 SETD2 Knockout cells and flow cytometry analysis

The coding sequence of NSD2 (amino acids 992–1240, UniProt No: O96028) WT and T1150A catalytic domains tagged with nuclear localization sequence (NLS) of the SV40 Large T-antigen were cloned into the mVenus-C1 plasmid backbone (provided by Steven Vogel, Addgene plasmids no. 27794) (Koushik et al., 2006) by Gibson-Assembly (NEB). NLS-NSD2-mVenus WT and T1150A plasmids were transfected into 70 % confluent HEK293 SETD2 knockout cells using FuGENE^®^ HD Transfection Reagent (Promega). NLS-mVenus empty vector was used as a negative control. After three days post transfection, cells were harvested and the transfection efficiency as well as expression of the NSD2 constructs in HEK293 cells were evaluated based on the mVenus reporter by flow cytometry (MACSQuant VYB, Miltenyi Biotec) and western blot using an anti GFP antibody (ThermoFisher, PA1-980A, 1:1000). Data analysis was performed using the FlowJo software (Treestar).

### 2.11 Cell lysis and immunoblotting

Both, parental and SETD2 KO HEK293 cells, were lysed for immunoblotting. For nuclear lysate enrichment, cell pellets were resuspended first in lysis buffer (10 mM Tris/HCl pH 8, 10 mM NaCl and 0.2% NP-40) supplemented with protease inhibitor cocktail (cOmplete ULTRA tablets, Mini, EDTA -free, EASYpack, Roche) and kept shaking on rotor for 30 min at 4 °C. Next, the samples were centrifuged at 4 °C and the supernatant was removed. A second, high salt lysis buffer (50 mM Tris/HCl pH 7.5, 1.5 mM MgCl2, 20% glycerol, 420 mM NaCl, 25 mM NaF and 1 mM Na3Vo4) supplemented with protease inhibitor cocktail was used to resuspend the nuclear pellets thoroughly with vortexing and sonification. Afterwards, the samples were spun down and the supernatant was aliquoted and flash frozen. Bradford assay was used to quantify the total protein amount of the lysate from the different samples and accordingly equal amounts of lysate were mixed with SDS-loading buffer and resolved by SDS-PAGE. Analysis was done by western blotting using the primary antibodies against H3K36me3 (ab9050, 1:2000), H3K36me2 (ab9049, 1:2500), H3 (ab1791, 1:10000) or GFP (ThermoFisher, PA1-980A, 1:1000) and as secondary antibody anti-rabbit HRP (Na934v, GE Healthcare, 1:5000). The signal was detected by chemiluminescence after the addition of Pierce™ ECL Western Blotting-substrate.

### 2.12 Steered molecular dynamics simulation (sMD)

Steered molecular dynamics simulations were performed in OpenMM 7.5.1 (Eastman and Pande, 2015; Eastman et al., 2017) utilizing the NVIDIA CUDA (https://docs.nvidia.com/cuda/) GPU platform. The systems were parameterized using the General Amber force field (GAFF) and AMBER 14 all atom force field (Case et al., 2014; Copeland et al., 2009; Wang et al., 2004) if not specified otherwise. The non-bonded interactions were treated with a cut-off at 10 Å. Additionally, the Particle Mesh Ewald method (Darden et al., 1993) was used to compute long range Coulomb interactions with a 10 Å nonbonded cut-off for the direct space interactions. Energy minimization of the system was performed until a 10 kJ/mol tolerance energy was reached. Simulations were run using a 2 fs integration time step. The Langevin integrator (Bussi and Parrinello, 2007) was used to maintain the system temperature at 300 K with a friction coefficient of 1 ps^-1^. The initial velocities were assigned randomly to each atom using a Maxwell-Boltzmann distribution at 300 K. A cubic water box with a 10 Å padding to the nearest solute atom was filled with water molecules using the tip4p-Ew model (Horn et al., 2004). Production runs were performed under periodic boundary conditions and trajectories were written every 10,000 steps (20 ps).

The structures of human NSD2 WT and NSD2 T1150A (amino acids 992-1221) were modelled based on the cryo-EM structure of NSD2 E1099K, T1150A in a nucleosome complex (PDB 7CRO) (Li et al., 2021). Missing amino acids and the reverting mutations of K1099E and A1150T (as appropriate) were modelled using PyMOD 3.0 (Janson and Paiardini, 2021). The missing part of the post-SET loop (1207-1221) in PDB 7CRO was modelled based on the SET domain of SETD2 (PDB: 5V21) (Zhang et al., 2017) using PyMOD 3.0, since no structure of NSD2 complexed with the H3K36 peptide and post-SET loop has been resolved. Subsequently, the histone tail of PDB 7CRO was replaced by the H3K36 peptide from PDB 5V21 and methionine 36 mutated to lysine. The target lysine 36 was then manually deprotonated as required for the S_N_2 mechanism (Poulin et al., 2016; Zhang and Bruice, 2008). Methyl groups were introduced at the lysine sidechain nitrogen using PyMOL (Schrödinger, 2015).

Parametrization of methylated lysine in the different methylation states was accomplished using AMBER 14 GAFF and ff14SB (Maier et al., 2015). AdoMet was modelled based on the coordinates of AdoHcy and parametrized using ANTECHAMBER from AmberTools (18.0) (Wang et al., 2001) and placed ^~^27 Å away from the AdoMet binding pocket.

The Zn^2+^ ions were modelled using the cationic dummy atom method (Oelschlaeger et al., 2003; Pang, 2001; Pang et al., 2000). Cysteines 1499, 1501, 1516, 1520, 1529, 1533, 1539, 1631, 1678, 1680, 1685 were treated as unprotonated to ensure proper Zn^2+^ binding (Cheng and Zhang, 2007). The protein charge was neutralized and an ionic strength of 0.1 M NaCl was applied, by adding 43 Na^+^ and 40 Cl^-^ ions.

To equilibrate the solvent, a 5 ns pressure coupled equilibration with Monte Carlo barostat (Faller and De Pablo, 2002) was performed at a pressure of 1 atm. The C-alpha (Cα) atoms of NSD2, the peptide and the AdoMet atoms were restrained with a force of 100 and 5 kJ/mol x Å^2^, respectively. The restraints were taken off successively, starting with the NSD2 Cα restraints, followed by a 5 ns equilibration with the peptide and AdoMet still being restrained. Subsequently, the AdoMet and peptide restraints were removed as well, followed by 0.1 ns equilibration with no restraints. A distant dependent force of 0.2 x distance of centroid 1 (lysine 36 side chain nitrogen and its attached two hydrogen atoms) and centroid 2 (AdoMet methyl group and its attached 3 hydrogen atoms) (kJ/mol)/Å^2^) was used to pull the centre of mass (COM) of centroid 1, towards the COM of centroid 2. The lysine hydrogen atoms are replaced with carbon atoms upon methyl group introduction. Two additional weaker forces were used to guide AdoMet into a proper binding position in the AdoMet binding pocket (force2: 0.1 x distance of centroid 3 (AdoMet atoms N1, C2, N3) and centroid 4 (NSD2 L1202 atoms N, Cα, C) (kJ/mol)/Å^2^); force3: 0.05 x distance of centroid 5 (AdoMet atoms N0, Cα, Cβ) and centroid 6 (NSD2 F1149 atoms Cα, C, O) (kJ/mol)/Å^2^)). For production, sMD simulations were conducted for 100 replicates à 35 ns (total simulation time 3.5 μs).

In order to define criteria describing a successful docking of AdoMet, the following geometric requirements for a transition state (TS)-like conformation were derived from the known S_N_2 geometry of methyl group transfer reaction (Schnee et al., 2022) (Supplementary Figure 12):

1. the distance between the lysine Nε and AdoMet methyl group C-atom is < 4 Å
2. the angle between the lysine Nε - lysine Cδ bond and the virtual bond between lysine Nε and the AdoMet methyl group C-atom is in a range of 109° ± 30°
3. the angle between the lysine Nε-AdoMet methyl group C-atom and AdoMet methyl group C-atom - AdoMet S-atom bonds is in a range of 180° ± 30°

Data analysis was performed utilizing MDTraj (1.9.4) (McGibbon et al., 2015) to calculate the distances and angles necessary for the geometric criteria of an S_N_2 TS-like conformation and the RMSD. All structures were visualized using PyMOL (2.4.1).

### 2.13 Volume estimation of the NSD2 active pocket

General simulation parameters and starting structures of NSD2 WT and T1150A complexed with the H3K36 peptide were modelled as described above for the sMD experiments. AdoMet was positioned in the AdoMet binding pocket based on the coordinates of AdoHcy in PDB 7CRO (Li et al., 2021). A 5 ns pressure coupled equilibration with Monte Carlo barostat (Faller and De Pablo, 2002) was performed at a pressure of 1 atm. NSD2 and peptide Cα atoms as well as cofactor AdoMet atoms were restrained with a force of 100 and 5 kJ/mol x Å^2^, respectively. The restraints were taken off successively, starting with the Cα restraints, followed by a 5 ns equilibration with only AdoMet restrained. Subsequently, the AdoMet restraints were removed as well followed by 5 ns equilibration with no restraints. For production, 30 replicates à 100 ns were performed (total simulation time 3 μs).

Analysis of the volumes around lysine 36 was performed using POVME3 (Wagner et al., 2017) with a grid spacing of 0.4 Å, a distance cut of 0.4 Å, a contiguous points criterion of 3 and a convex hull exclusion. The coordinates of the inclusion spheres are: 35.30, 40.30, 34.55 and 30.00, 39.77, 34.69 each with a radius of 3.0 Å (Supplementary Figure 13). Out of the total simulation time of 3 μs, 10 % of the simulation frames were randomly chosen and the volume calculated. This process was done in triplicates for each methylation state (me0, me1, me2) for NSD2 WT and NSD2 T1150A.

### 2.14 Differential gene expression analysis of NSD2 WT and T1150A

The Cancer cell line encyclopedia (CCLE) (https://sites.broadinstitute.org/ccle) and Catalogue of somatic mutants in cancer (COSMIC) (cancer.sanger.ac.uk) databases were screened for hematological cancer cell lines harboring the NSD2 T1150A mutant, which revealed a diffuse large B cell lymphoma cell line (OCILY-18) containing the mutant of interest. Four more NSD2 WT control cell lines were selected (OCILY-1, OCILY-7, OCILY-10 and DOHH2) for comparison with OCILY-18 (Barretina et al., 2012; Klijn et al., 2015; Rouillard et al., 2016; Tate et al., 2019). The control cell lines were selected to have the same cancer disease subtype (diffuse large B cell lymphoma) as NSD2 T1150A cancer cell line OCILY-18 and at the same time not to carry other mutations in NSD1, NSD2, NSD3 and SETD2. Additionally, any NSD2 WT containing cell line showed overexpression in NSD enzymes in comparison to other control cell lines was not selected. Moreover, gene expression data and mutational profiles of the selected cell lines needed to be available. The GSE57083 dataset, which contains the RNA expression microarray data of all mutant and control cell lines in the same platform (GPL570), was used to retrieve the differentially expressed genes (Log2 fold change (FC) ≥2 or ≤-2, adjusted p-value < 0.05). The analysis was done using the GEO2R analysis tool (https://www.ncbi.nlm.nih.gov/geo/geo2r/) (Barrett et al., 2013) with implemented Benjamini & Hochberg correction (False discovery rate) to obtain adjusted p-value and correction for multiple testing. The probes which are not specifically assigned to a single gene were removed.

The differentially upregulated and downregulated genes were analyzed for significantly enriched gene ontology (GO) terms and biological processes using the (Enrichr) analysis tool (https://maayanlab.cloud/Enrichr/) at FDR <0.05 using Benjamini & Hochberg correction (Chen et al., 2013; Kuleshov et al., 2016; Xie et al., 2021). In order to investigate the most enriched histone modification(s) and transcription factors (TFs) at the differentially expressed genes, the ChIP-seq data in the ENCODE-histone modifications and ENCODE-TFs databases were analyzed using the Enrichr analysis tool (Chen et al., 2013; Kuleshov et al., 2016; Xie et al., 2021). All hits were ranked according to their adjusted p-value (significance when adjusted p-value <0.05 using Benjamini & Hochberg correction).

### 2.15 Statistics

T-Tests were conducted with Excel using the specified settings. P-values based on binomial distributions were calculated with Excel using the Binom.dist function.

## 3 Results and discussion

### 3.1 Selection of NSD2 and NSD1 somatic cancer mutants to study

For the selection of somatic cancer mutants in NSD2 that potentially affect the methyltransferase activity, we screened the Catalogue of somatic mutants in cancer (COSMIC) database and filtered for missense mutations in the catalytic SET domain (Figure 1A). The most frequently observed NSD2 SET domain missense mutation is E1099K, which is followed by T1150A. Both mutations were specifically observed in patients with hematological cancers. While NSD2 E1099K has already been well characterized (Jaffe et al., 2013; Li et al., 2021; Oyer et al., 2014; Pierro et al., 2020; Sato et al., 2021; Swaroop et al., 2019), very little was known about T1150A when we started our work, which is why NSD2 T1150A was selected as the focus of this study. In addition, we were interested to compare the NDS2 data with one of its homologues, NSD1, and screened the COSMIC profile of the NSD1 SET domain as well (Figure 1B). Indeed, the NSD1 T2029A mutation, corresponding to NSD2 T1150A, has been observed in hematological malignancies as well. Moreover, the NSD1 COSMIC profile contains additional missense mutations and the Y1971C and R2017Q/L mutations have similar frequencies as T2029A. They were observed in different types of solid cancers like digestive tract, endometrial and other soft tissues carcinomas, and we included them in our study as well.

**Figure 1:**
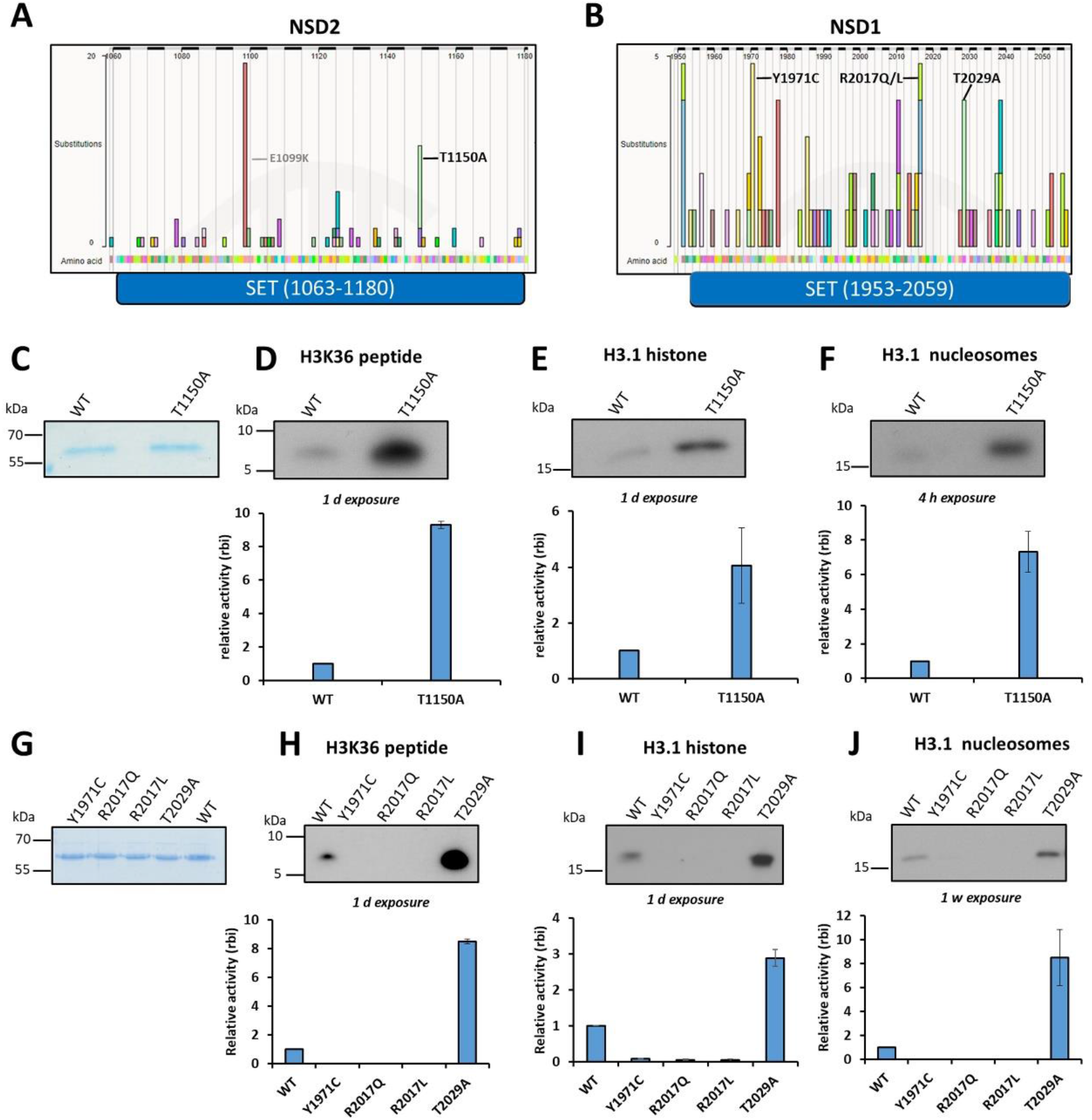
Selection and methyltransferase activity analysis of NSD2 and NSD1 missense cancer mutants. (A-B) Somatic cancer missense mutations in the SET domains of NSD2 (A) and NSD1 (B). The selected mutations investigated in this study are labelled in black while the previously investigated NSD2 E1099K mutation is labelled in gray. (C, G) Coomassie BB stained SDS-polyacrylamide gels depicting equal amounts of purified GST-tagged WT and mutant proteins for NSD2 (C) and NSD1 (G). (D-F, H-J) Methyltransferase assays of NSD2 (D-F) and NSD1 (H-J) cancer mutants. The methyltransferase assays were conducted using either H3K36 peptide (D, H), H3.1 recombinant protein (E, I), or H3.1 recombinant mononucleosomes (F, J) as methylation substrate and radioactively labelled AdoMet. Autoradiographic pictures of the methyltransferase assay are shown in the upper panels and the corresponding quantitative analysis in the lower panels. The mutant activities are displayed relative to WT enzymes based on the relative band intensity (rbi). The data are expressed as means ± SEM for at least 2 independent replicates.

### 3.2 Activity analysis of the somatic cancer mutants of NSD2 and NSD1

First, we wanted to compare the methyltransferase activity of the mutants with the corresponding wildtype (WT) enzymes using in vitro methyltransferase assays. To this end, the GST-NSD2-SET WT and the corresponding T1150A mutant, created by site-directed mutagenesis, were overexpressed in *E. coli* and purified from bacterial cells (Figure 1C). The catalytic SET domains of NSD1 and NSD2 were shown before to recapitulate the full length enzyme in activity as well as substrate and product specificity (Li et al., 2009; Poulin et al., 2016; Qiao et al., 2011). Equal concentrations of both GST-NSD2 WT and T1150A mutant were mixed in methylation buffer supplemented with radioactively labelled AdoMet with one of 3 different histone substrates, H3K36 unmodified peptide, H3.1 recombinant protein or recombinant H3.1 mononucleosomes. The methylation signal was detected afterwards by autoradiography (Figures 1D-F, Supplementary Figure 1). Consistently, the T1150A cancer mutant showed hyperactivity compared to WT on all three different histone substrates. Quantification of the autoradiographic signals revealed an around 9-fold higher activity on the H3 peptide substrate, 4-fold more on recombinant H3.1 protein and 7-fold on recombinant mononucleosomal substrates (Figures 1D-F). This finding is in agreement with a recent study which also reported hyperactivity of NSD2 T1150A (Sato et al., 2021) and the observation that overexpression of NSD1 and NSD2 and other activating mutations like NSD2 E1099K are frequently observed in blood cancers as described above. Additionally, these results suggest that the NSD2 T1150A mutant mediates carcinogenesis through a gain-of-function mechanism.

For comparison with NSD1, we used an existing murine NSD1-SET domain WT construct (Kudithipudi et al., 2014b) that shares >95% identity with the human protein. We then cloned and purified the NSD1 mutants (Figure 1G) and tested their methyltransferase activity as well. Consistently, NSD1 T2029A also showed an enhanced activity compared to NSD1 WT on the different histone substrates (8.5-fold for peptide and nucleosomal substrates, and 3-fold on the protein substrate) (Figures 1H-J). In contrast, the other NSD1 cancer mutants (Y1971C, R2017Q and R2017L), which are mostly observed in different types of solid tumors, abolished the NSD1 activity (Figures 1H-J). The R2017Q results agree with a previous study investigating this mutant in the context of SOTOS syndrome (Qiao et al., 2011). The difference in the activity of the different NSD1 cancer mutants ranging from inactivity to hyperactivity suggests that NSD1 can act as oncogene or tumor suppressor in a tumor type specific manner. The NSD1 loss-of-function effect of the three cancer missense mutations discovered here is in agreement with reports describing epigenetic silencing of NSD1 by promotor hypermethylation or genetic deletions in solid tumor cell lines and cancer patients (Berdasco et al., 2009; Brennan et al., 2017; Network, 2015; Su et al., 2017). These NSD1 silenced tumors show decreased intergenic H3K36me2 and DNA hypomethylation of CpG islands and intergenic regions at affected genomic loci (Su et al., 2017; Tauchmann and Schwaller, 2021). Mechanistically, the effects of the Y1971C and R2017Q/L mutations can be explained in the light of the recently published cryo-EM structure of NSD2 in complex with a nucleosome containing H3K36M. This structure shows that NSD2 Y1092 (which corresponds to NSD1 Y1971) contacts the H3K36M side chain together with two more aromatic residues (F1177 and Y1179) and an additional hydrophobic one (L1120) (Sato et al., 2021). The Y1971C mutation disrupts this aromatic cage which is needed for positioning and deprotonation of the target lysine explaining the loss of catalytic activity caused by the mutation. In an NSD1-AdoMet structure, R2017 is observed to bind to three aromatic amino acids, Tyr1870, Tyr1997, and Phe2018 through H-bonds and cation-π interactions (Qiao et al., 2011). These residues are known to be essential for the activity of many SET domain containing PKMTs (Qiao et al., 2011). Moreover, R1138, the NSD2 residue corresponding to NSD1 R2017, is involved in (C-H-O) hydrogen bonding between its main chain carbonyl oxygen and the C-atom of the AdoMet methyl group suggesting that it has a direct role in catalysis (Poulin et al., 2016). Collectively, these findings can explain that the mutation of R2017 to leucine or glutamine abolishes the enzymatic activity of NSD1.

To investigate if there are larger differences in folding between the NSD1/2 WT enzymes and their corresponding cancer mutants, CD spectroscopy was conducted with all proteins. All spectra were similar indicating comparable secondary structure and folding between WT enzymes and their corresponding cancer mutants (Supplementary Figure 2). Collectively, these results confirm that NSD2 T1150 and its equivalent residue in NSD1, T2029, are important for controlling the methyltransferase enzymatic activity and their mutation to alanine increases the activity of both enzymes. In contrast, NSD1 residues Y1971 and R2017 are essential for enzymatic activity and the Y1971C and R2017Q/L exchanges cause a loss-of-function.

### 3.3 T1150A/T2029A change the enzyme product specificity on protein and nucleosomal substrates

Since only NSD2 T1150A and its homologue NSD1 T2029A exhibited catalytic activity, we decided to characterize these hyperactive mutants in more detail. NSD1 and NSD2 are both known to generate H3K36 methylation only up to the dimethylation state in vitro and in vivo (Husmann and Gozani, 2019; Li et al., 2019; Li et al., 2009; Qiao et al., 2011), while SETD2 is the sole human enzyme to catalyze up to trimethylation of H3K36 in somatic cells (Edmunds et al., 2008). The change in H3K36 methylation state can result in different biological responses (DiFiore et al., 2020; Li et al., 2019; Wagner and Carpenter, 2012), but the exact molecular reason for the dimethyl product specificity of NSD1 and NSD2 is unknown. We were interested to investigate whether the hyperactive NSD2 T1150A cancer mutant can change the NSD2 product specificity to also generate H3K36me3. To test this hypothesis, equal molar concentration of NSD2 WT and T1150A were mixed with recombinant H3.1 protein in methylation buffer supplemented with unlabeled AdoMet. After methylation, the samples were loaded onto an SDS-polyacrylamide gel and the methylation state of methylated histone proteins was analyzed by western blot using antibodies specific for either H3K36me2 or H3K36me3. Strikingly, while both NSD2 WT and T1150A mutant could catalyze the histone H3K36 dimethylation, only the NSD2 T1150A mutant was able to generate H3K36me3 (Figures 2A-B). We confirmed the specificity of the H3K36me3 specific antibody using SPOT peptide array binding experiments (Supplementary Figure 4A). Since the H3.1 recombinant mononucleosomes are a more physiologically relevant substrate, they were used in a similar methyltransferase assay. Again, it was observed that the NSD2 T1150A variant, but not NSD2 WT, catalyzed trimethylation of H3.1 nucleosomes (Figure 2C, Supplementary Figure 3A). Noteworthy, the same effect was also observed using the NSD1 T2029A mutant on both the H3.1 protein and mononucleosomal substrates (Figures 2D-F, Supplementary Figure 3B). Moreover, and in order to rule out any possible cross reactivity of the H3K36me3 specific antibody with H3K36me2, the methyltransferase assay was repeated with titrating the amount of the hyperactive NSD1 T2029A mutant until equal dimethylation activity was achieved in comparison with NSD1 WT (Supplementary Figure 4B). At such conditions, the H3K36me3 methylation signal was only detected with the T2029A mutant (Supplementary Figure 4C). This confirms the specificity of H3K36me3 antibody and indicates that the detected H3K36me3 signal is indeed due to trimethylated H3K36 introduced by the T2029A mutant and not to H3K36me2 cross reactivity. As H3K36me3 is observed under conditions of similar H3K36me2 activity of WT and T2029A, the H3K36me3 generation by T2029A is not a simple consequence of its hyperactivity, but due to a real change of product specificity of the mutant.

**Figure 2:**
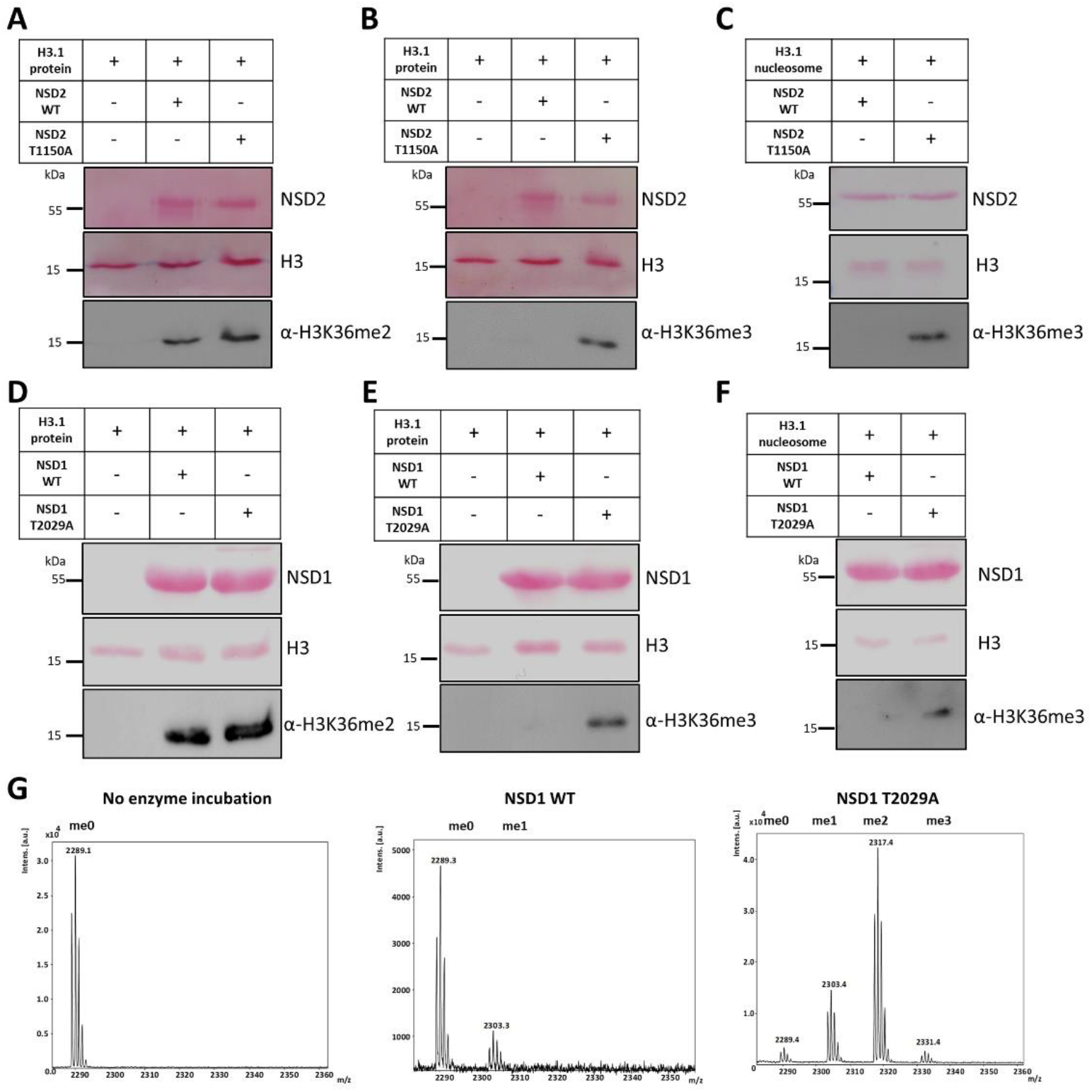
Product specificity change of NSD2 T1150A and NSD1 T2029A compared to WT enzymes on H3.1 protein and nucleosomes in vitro. (A-B) Western blot analysis using H3K36me3 specific antibody (A) or H3K36me2 specific antibody (B) after methylation of H3.1 recombinant protein with NSD2 WT or T1150A. (C) Western blot analysis using the H3K36me3 specific antibody after methylation of H3.1 nucleosomes with NSD2 WT or T1150A. (D-E) Western blot analysis signals using H3K36me3 specific antibody (D) or H3K36me2 specific antibody (E) after methylation of H3.1 recombinant protein with NSD1 WT versus T2029A mutant. (F) Western blot analysis using the H3K36me3 specific antibody after methylation of H3.1 nucleosomes with NSD1 WT or T2029A. In all panels, the equal loading of substrates and enzymes is shown by Ponceau S staining. (G) Methylation of the H3K36me0 peptide by NSD1 WT and T2029A analyzed by MALDI mass spectrometry. The figures show from left to right the mass spectra of the H3K36 peptide without NSD1 enzyme incubation, the H3K36 peptide after incubation with NSD1 WT or NSD1 T2029A. The samples were incubated in methylation buffer containing AdoMet for 4 h at 37 °C. The masses of the corresponding peptides are 2289 Da (H3K36me0), 2303 Da (H3K36me1), 2317 Da (H3K36me2) and 2331 Da (H3K36me3).

### 3.4 Mass spectrometry analysis revealed the trimethylation activity of NSD1 T2029A on an H3K36 peptide substrate

In order to confirm the trimethylation activity of the hyperactive cancer mutants, we applied mass spectrometry which is independent of the use of H3K36me3 specific antibodies. Due to the lower sensitivity of the mass spectroscopy coupled methylation assays as compared with radioactive or antibody based methods, sufficient catalytic activity was only observed with the purified NSD1 proteins. Accordingly, NSD1 WT or T2029A mutant were used in equal molar concentration in a methyltransferase assay using the unmodified H3(26-44) peptide as substrate. After the reaction, the methylation states of histone peptides were identified by MALDI-TOF mass spectrometry. As shown in Figure 2G, T2029A was able to methylate the H3K36 peptide resulting in all methylation states (mono-, di- and trimethylation) which is in agreement with our western blot results. In contrast, only monomethylated H3K36 peptide was detected in the reaction with NSD1 WT.

Additionally, the H1.5 K168 peptide was reported previously to be an NSD1 histone target with even stronger methylation than H3K36 peptide in vitro (Kudithipudi et al., 2014b). We, therefore, tested if NSD1 T2029A mutant can catalyze trimethylation on the H1.5 K168 peptide as well. First the methyltransferase activity of NSD1 T2029A on H1.5(160-176) peptide was compared to NSD1 WT enzyme in radioactive methyltransferase assay and the signal was detected by autoradiography (Supplementary Figure 5A). As observed previously with the H3K36 substrate, T2029A was hyperactive relative to NSD1 WT (Supplementary Figure 5A). Interestingly and in agreement with H3K36 trimethylation activity of T2029A, in methylation assays coupled to mass spectrometry readout, trimethylation of the H1.5 K168 peptide was observed with T2029A but not NSD1 WT (Supplementary Figure 5B-D). In summary, these results document a unique trimethylation activity of the NSD2 T1150A and NSD1 T2029A cancer mutants. This activity could rewrite the H3K36 methylation landscape leading to massive biological responses which may explain the carcinogenic effect of these mutations.

### 3.5 NSD2 T1150A shows enhanced complex formation with H3K36me2 and AdoMet compared to NSD2 WT

We next aimed to characterize the mechanism behind this exceptional trimethylation catalysis of the NSD2 T1150A mutant on H3K36 using MD simulations. For the correct setup of these experiments, information about the reaction mechanism was essential, in particular whether the enzyme acts in a processive or distributive manner. Many SET-domain PKMTs were shown to exhibit a processive mode of action for the stepwise methylation of their target lysine residues (Dirk et al., 2007; Kwon et al., 2003; Patnaik et al., 2004; Zhang et al., 2003). This means that after one methyl transfer reaction, the cofactor product S-adenosyl-L-homocysteine (AdoHcy) dissociates and a new AdoMet molecule is bound without dissociation of the peptide substrate. To confirm this mechanism for the NSD enzymes, we conducted methylation reactions of the H3K36me1 peptide with NSD1 T2029A (the only enzyme that was active enough for this kind of analysis) and detected the product methylation states by MALDI mass spectrometry. Noteworthy, starting with H3K36me0 a clear peak of dimethylated H3K36 product and also some trimethylation was observed as mentioned before (Figure 2G). In contrast, much lower methylation was detectable when the reaction was started under the same conditions with the H3K36me1 substrate (Supplementary Figure 6B). This result is not compatible with a distributive reaction mechanism.

In order to investigate the mechanistic basis of the difference in product specificity between NSD2 WT and T1150A, steered molecular dynamics (sMD) simulations were applied. In the sMD approach, external forces are used to guide bimolecular association processes (Yang et al., 2019). Thereby, reactions that otherwise would be too slow to be modelled in MD simulations are accelerated and at the same time the conformational sampling is concentrated along a specific, predefined reaction coordinate. In a processive reaction mechanism, higher product methylation states are achieved by the release of AdoHcy after one methylation reaction followed by the binding of a new AdoMet to the enzyme-peptide complex. Consequently, the AdoMet association process to enzyme-peptide complexes already containing H3K36me1 or me2 peptides was modelled to study the potential generation of higher H3K36 methylation states up to H3K36me3. For this, NSD2-peptide complexes with the H3K36me1 and me2 peptide bound in the NSD2 peptide binding cleft and devoid of AdoMet were modelled using the cryo-EM structure of NSD2 E1099K, T1150A bound to a nucleosome and the SET domain of SETD2 complexed with H3K36M as templates (PDB 7CRO and 5V21, see also the method description and Supplementary data for details and structures). To simulate the AdoMet association process into the active site of the NSD2-peptide complexes, it was placed 27 Å above its binding pocket and a weak attractive force of 0.2 kJ/(mol x Å^2^) was applied between the Nε-atom of lysine 36 and the methyl group C-atom of AdoMet (Figure 3A). In order to define criteria describing a successful docking of AdoMet, the geometric requirements for a transition state (TS)-like conformation were applied, which were derived from the known S_N_2 geometry of methyl group transfer reactions (Schnee et al., 2022).

**Figure 3:**
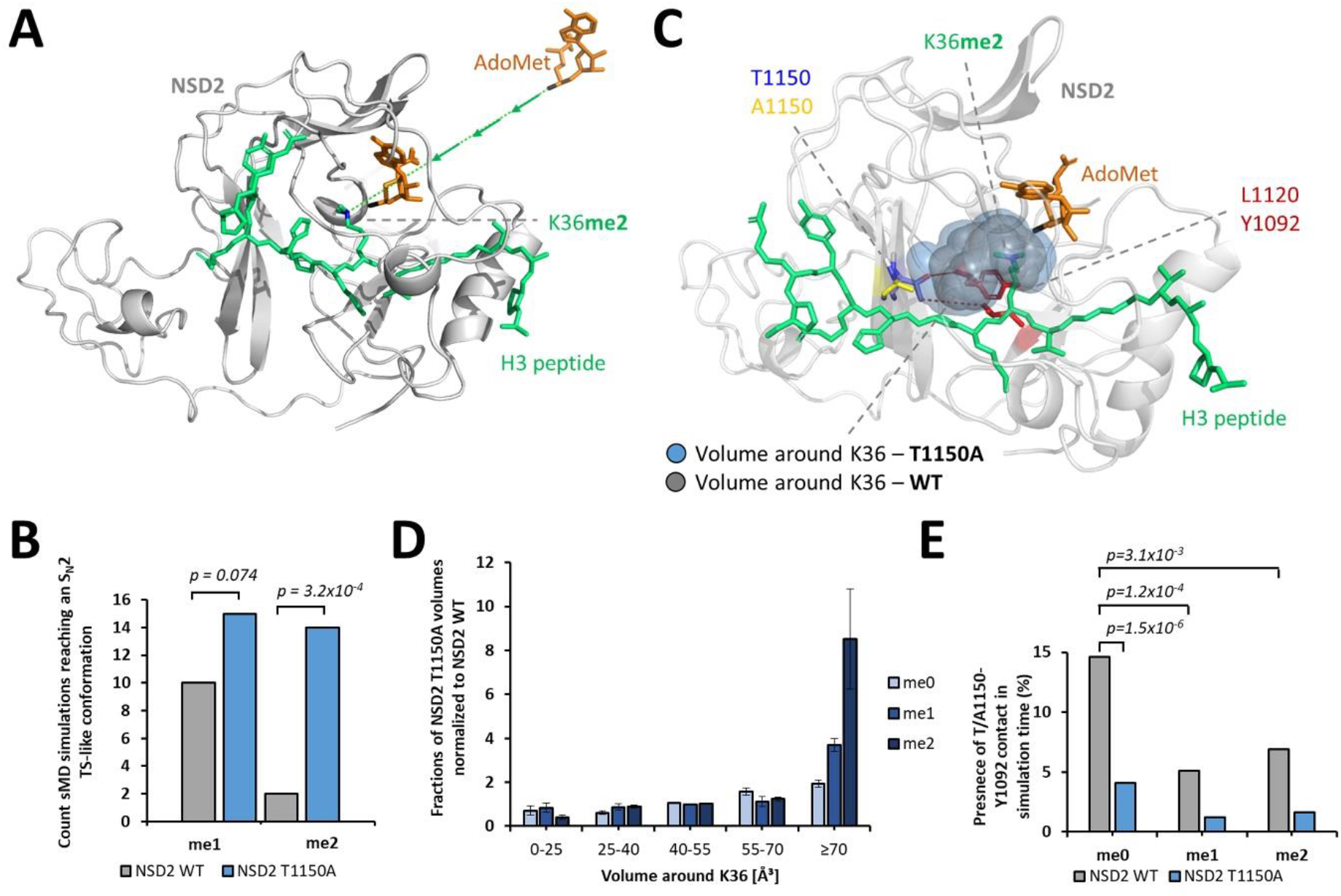
sMD simulation of AdoMet association to WT and T1150A NSD2 – peptide complexes and measurement of the active site pocket volume in NSD2 WT and T1150A complexes. (A) Starting position of the sMD simulation replicates, in which H3K36me1 or me2 are bound to NSD2 and AdoMet was positioned 27 Å away from the AdoMet binding pocket. (B) Number of successful docking events reaching a TS-like conformation in 100 sMD simulations à 35 ns with H3K36me1 or H3K36me2 peptides complexes (Supplementary Figure 12). (C) Structure of the complex of NSD2 with bound H3K36me2 and AdoMet as well as the corresponding volumes around K36 for NSD2 WT and T1150A. Red lines indicate the contacts of T1150 with Y1092 and L1120. (D) Distribution of the volumes around K36 in Å^3^ observed in MD simulations of NSD2 T1150A – peptide – AdoMet complexes, normalized to the corresponding values for NSD2 WT (for unnormalized data see Supplementary Figure 13). (E) Presence of the contact between T1150 and Y1092 in % of the simulation time in 30 simulations à 100 ns. Corresponding p-values were determined by two-tailed T-Test assuming equal variance on the basis of the 30 sMD replicates.

One hundred sMD simulations à 35 ns were performed for each NSD2 WT and T1150A and frames were recorded every 20 ps. Then, the number of docking events were monitored which fulfilled the success criteria in at least one frame. The analysis of the sMD simulations revealed that both proteins were able to establish S_N_2 TS-like conformations, however with significant differences. With the H3K36me1 substrate, the analysis revealed 10 successful docking simulations for NSD2 WT and 15 for NSD2 T1150A corresponding to a non-significant difference (Figure 3B). In contrast, with the H3K36me2 substrate, NSD2 WT accommodated AdoMet successfully into the binding pocket in only 2 out of 100 simulations, whereas NSD2 T1150A did this in 14 out of 100 simulations (p-value 3.2 x 10^-4^) (Figure 3B). An analysis of the RMSD of successful AdoMet association compared to already bound AdoMet showed that the positions were in good agreement (Supplementary Figure 7A and B). An example of a successful docking event is provided in Supplementary Movie 1. In conclusion this sMD experiment fully recapitulates the biochemical data described above. It identified an enhanced capability of NSD2 T1150A to accommodate AdoMet when H3K36me2 is already bound as compared to NSD2 WT which can explain its ability to generate H3K36me3.

### 3.6 NSD2 T1150A has an increased active site volume to accommodate H3K36me2 and AdoMet

The sMD experiments showed an enhanced ability of NSD2 T1150A to bind H3K36me2 and AdoMet simultaneously compared to NSD2 WT. To analyze the molecular mechanism of this effect, the volume of the active site pocket was measured during MD simulations and the contacts of the active site amino acids were examined. Firstly, an analysis of the active site volume was carried out by simulating NSD2 complexed with AdoMet and the H3K36 peptides in different methylation states. By having AdoMet already bound, a standardized comparison between NSD2 WT and NSD2 T1150A can be made and undersampling of NSD2 WT frames with bound AdoMet was avoided. For this, 30 MD simulations à 100 ns were conducted for each NSD2 WT and T1150A. Out of this pool, 5000 randomly selected snapshots were used to calculate the active site volume around K36 for each protein complexed with H3K36me0, me1 or me2 (Figure 3C). The analysis of the calculated volumes shows that large volumes (≥70 Å^3^) occur more frequently for NSD2 T1150A and lower volumes (0-25 Å^3^) more frequently for NSD2 WT. This effect increases with higher methylation levels of K36. For K36me2, large volumes occur 8.5-fold more often for NSD2 T1150A compared to NSD2 WT (p-value 0.015, calculated by two-tailed T-Test assuming equal variance based on three replicates of the analysis) (Figure 3D). The strong elevation of this effect with higher H3K36 methylation levels suggests that the active site tends to collapse with higher methylation levels and this effect is more pronounced with WT than with the mutant. Overall, these findings clearly explain the increased capability of T1150A to accommodate AdoMet and H3K36me2 simultaneously.

Since the T1150A mutation is the only difference between the two proteins, the increased volume of the active site pocket must be a direct consequence of the mutation. MD simulations of modelled complexes of NSD2 with the H3K36me0, me1 and me2 peptides revealed contacts of T1150 to Y1092 and L1120 (Figure 3C, Supplementary Figure 7C). Contacts were considered as established if the distance of a pair of heavy atoms from both amino acids was below 4.5 Å. An H-bond between the hydroxyl group of T1150 and the backbone amide of Y1092 was established in 15% of the simulation time (30 replicates of 100 ns) with H3K36me0, but only 5% and 7% in the case of me1 and me2 (Figure 3E and Supplementary Data). This contact orients Y1092 and restricts the volume of the active site pocket consequently disfavoring the interaction with me1 and me2 substrates. In case of A1150, the contact to Y1092 is much less frequent, which supports the further methylation of me1 and me2 substrates. The hydrophobic contact between the T1150 side chain methyl group and the Cδ-atoms of L1120 was observed in 53% of the simulation time (average of me0, me1 and me2), but only in 24% of the time in case of T1150A (p-value 1.2×10^-14^, calculated by two-tailed T-Test assuming equal variance based on 30 replicates MD analysis) (Supplementary Data). This interaction orients L1120 and also restricts the volume of the active site pocket. Hence, the disruption of contacts to Y1092 and L1120 is likely the reason for the enlargement of the active site volume of the T1150A mutant leading to its activity as trimethyltransferase.

As mentioned previously, Y1092 and L1120 surround the H3K36 side chain and the mutation of NSD1 Y1971 (corresponding to NSD2 Y1092) to cysteine abolished the enzyme activity. The concept of the size of the enzyme active site pocket as a limiting factor controlling the PKMT product specificity has been developed by comparing the Rubisco large subunit methyltransferase LSMT, a lysine trimethyltransferase, with the monomethyltransferase SET7/9 (Trievel et al., 2003). It had been further refined by showing that a Tyr/Phe switch in the lysine binding pocket can control the PKMT product specificity, where tyrosine favors the smaller sized product (mono- and dimethylation) while phenylalanine switches the specificity to di- and trimethylation (Collins et al., 2005). This concept was experimentally validated for several PKMTs (Collins et al., 2005; Couture et al., 2008; DiFiore et al., 2020; Guo and Guo, 2007; Weirich et al., 2015; Wu et al., 2010) and also rationalized by MD and QM simulations (Zhang and Bruice, 2008). Moreover, the G9a residue Y1067, corresponding to NSD2 Y1092, has been shown to control the dimethylation product specificity of G9a and its mutation to phenylalanine converted G9a to a trimethyltransferase (Wu et al., 2010). The loss of the positioning of NSD2 Y1092 in the T1150A mutant may have a similar effect, which could explain the switch from dimethyltransferase to trimethyltransferase activity in the T1150A mutant. Moreover, NSD2 L1120 is the only amino acid residue in close proximity to the H3K36 side chain which is not conserved between the dimethyltransferase NSD2 and the trimethyltransferase SETD2. SETD2 contains a less bulky methionine at this place suggesting that L1120 might also have an important role in controlling the product specificity of NSD2 (Figure 5A). The loss of the contact of T1150 to L1120 in the T1150A mutant might increase the flexibility of this residue and thereby destroy its control of the NSD2’s product specificity. In summary, our findings unraveled key mechanistic features controlling the product specificity of this class of enzymes by the accurate positioning of residues surrounding the substrate H3K36 side chain.

**Figure 4:**
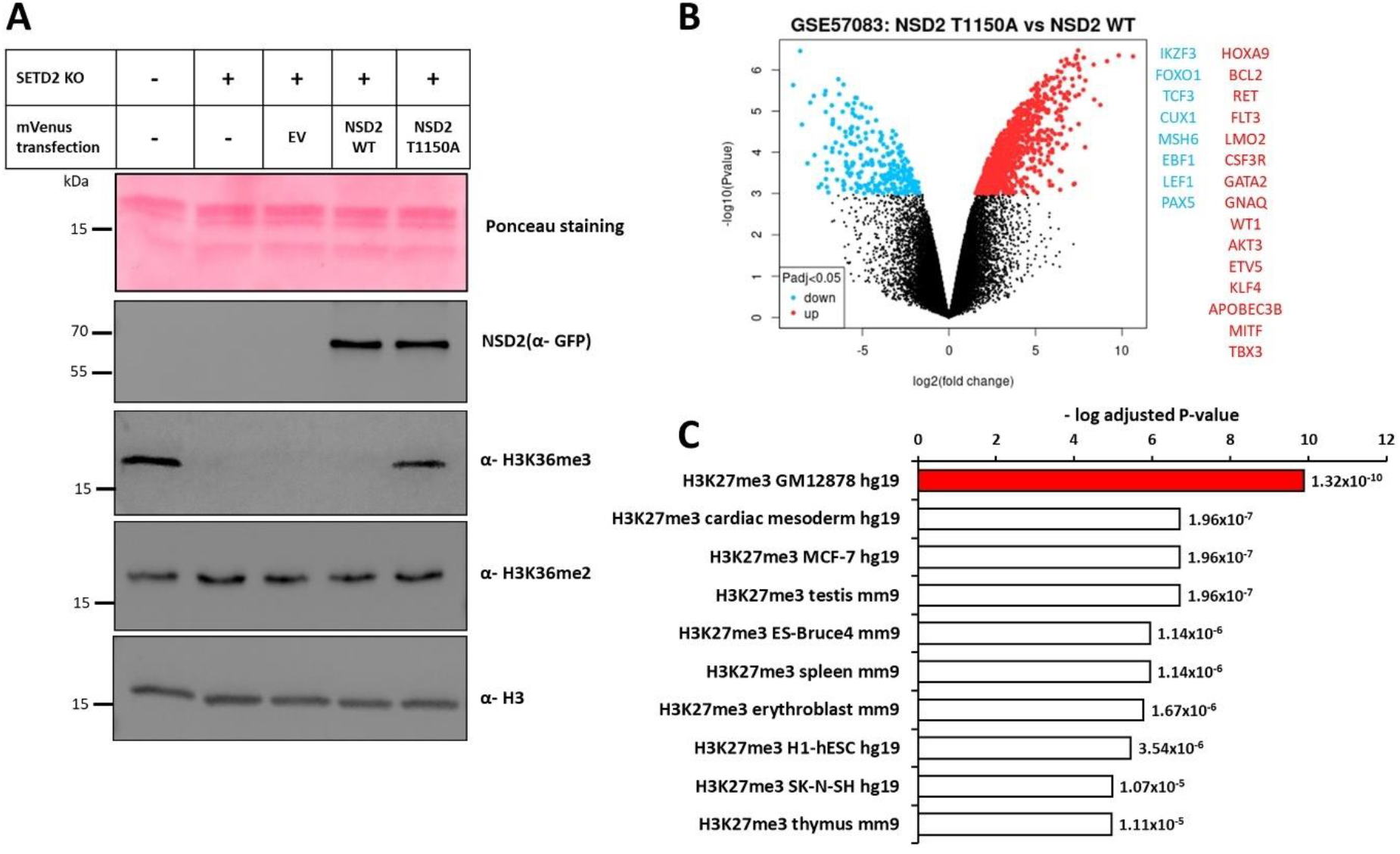
NSD2 T1150A introduces H3K36me3 in human cells and leads to upregulation of genes by competition with H3K27me3. (A) Immunoblotting of cell lysate of parental HEK293 cells, SETD2 KO HEK293 cells and SETD2 KO HEK293 cells transfected with different mVenus constructs (EV, NSD2 WT, NSD2 T1150A). (B) Volcano plot showing the differential gene expression analysis between the B-cell lymphoma cancer cell line (OCILY-18) harboring NSD2 T1150A versus 4 B-cell lymphoma cancer cell lines (OCILY-1, OCILY-7, OCILY-10 and DOHH2) which were selected not to contain mutations in NSD1, NSD2, NSD3 and SETD2 or overexpression of NSD enzymes. The significantly up- and downregulated genes are colored red and blue, respectively. Each dot is representing one gene probe corresponding in total to 504 significantly upregulated genes and 163 significantly downregulated genes (significant at adjusted p-value <0.05). Non-significant probes are colored black. The analysis was conducted using the GEO2R tool and GEO57083 expression array data. On the right side, examples of upregulated oncogenes (red) and downregulated TSGs (blue) mapped to COSMIC cancer gene census list are provided. (C) Analysis of the enrichment of histone modification(s) at the differentially upregulated genes shown in panel B (Log_2_ FC ≥2, adjusted p-value <0.05). The analysis was performed using the Encode database with the Enrichr analysis tool. All hits were ordered according to their adjusted p-values as indicated beside each bar. The bar diagram shows H3K27me3 in the B-lymphocyte cell line GM12878 as the most significant hit (marked in red).

**Figure 5:**
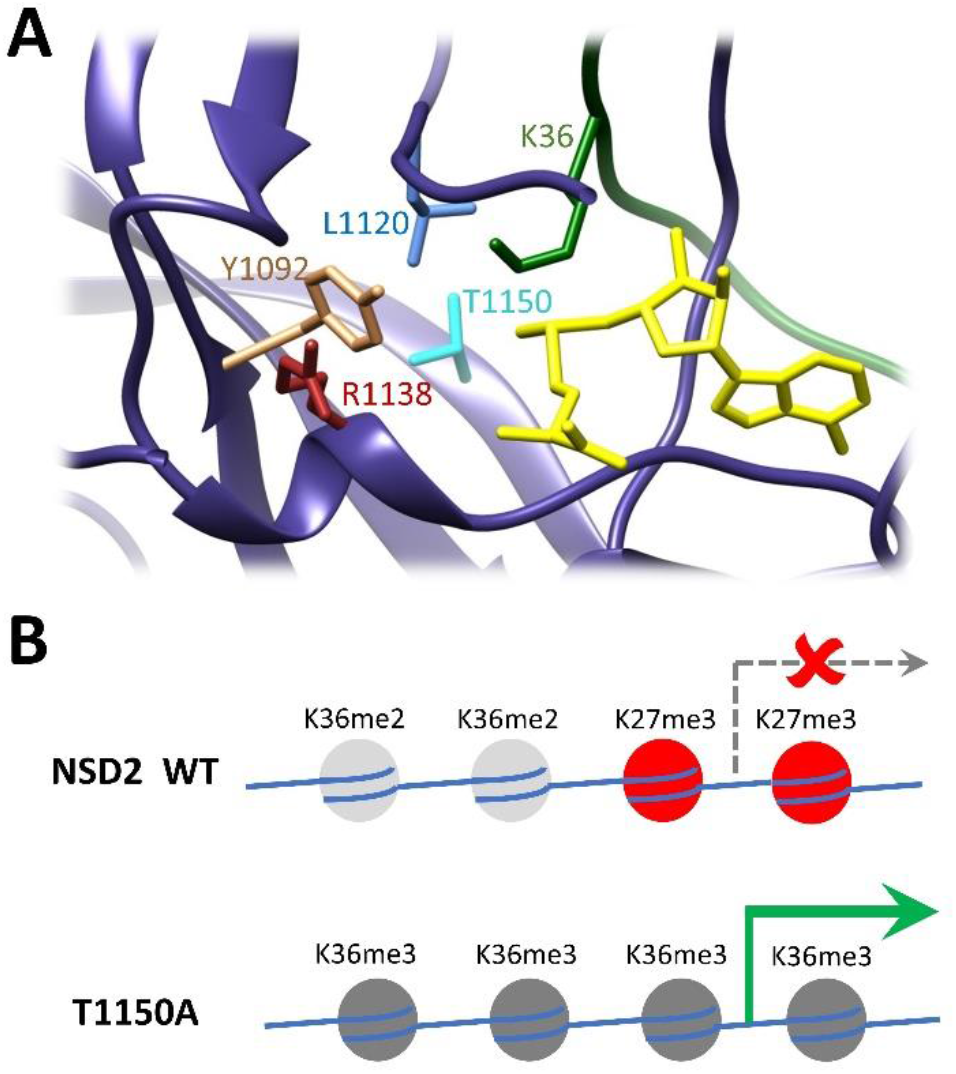
NSD2 structure showing the amino acids studied in this work and model for the biological effect of T1150A. (A) Example of one NSD2 structure in complex with AdoMet (yellow) and H3K36me0 peptide (green). NSD2 is shown in dark blue ribbon with atoms displayed for T1150, Y1092, L1120 and R1138. (B) Schematic model illustrating the biological effect of NSD2 T1150A cancer mutant. Aberrant deposition of K36me3 can lead to reduction in H3K27me3 followed by gene activation.

### 3.7 NSD2 T1150A catalyzes trimethylation of H3K36 in human cells

Next, we asked whether the NSD2 T1150A mutant can catalyze H3K36me3 in human cells as well and most importantly if this trimethylation will be due to direct catalysis by NSD2 T1150A and not mediated indirectly by the SETD2 enzyme (the sole human enzyme responsible for H3K36me3 deposition). To address this question, we first created a SETD2 knockout HEK293 cell line using CRISPER-Cas9 genetic engineering. Following literature data (Edmunds et al., 2008), the SETD2 KO was validated in the derived clonal cell lines by the absence of genomic H3K36me3 compared to parental cells (Supplementary Figure 8A). Analysis of genomic DNA revealed mutations in both alleles of the SETD2 gene in one selected clone (Supplementary Figure 8B and C). This step was followed by the expression of either NSD2 T1150A or NSD2 WT catalytic domain in the SETD2 KO cells and investigation of genomic H3K36me3 levels. To this end, NLS-mVenus-tagged NSD2 WT or NSD2 T1150A cancer mutant were transfected into the SETD2 KO cells together with NLS-mVenus empty vector as negative control. Transfection efficiency detected by the mVenus reporter fluorescence was analyzed by flow cytometry showing equal transfection yields (Supplementary Figure 9). Moreover, the equal expression of mVenus-tagged NSD2 WT or NSD2 T1150A was further confirmed by Western Blot using anti-GFP antibody (Figure 4A). Intriguingly, the overexpression of NSD2 T1150A in the SETD2 KO cell line led to a defined rise in genomic H3K36me3 levels, while cells overexpressing WT NSD2 did not show an increase of H3K36me3 when compared with untransfected cells or cells transfected with mVenus empty vector (Figure 4A). These results were confirmed in three independent transfection series. They directly demonstrate the capability of the NSD2 T1150A SET domain to generate H3K36me3 in human cells independently of SETD2. This result strongly suggests that the corresponding full-length NSD2 T1150A mutant has the same ability, as so far regions outside of the catalytic part of SET-domain PKMTs have not been shown to affect product methylation levels.

The aberrant trimethylation catalysis of NSD2 T1150A in cells can change the H3K36 methylation state which is known to be associated with diverse biological outcomes, because H3K36me2 and H3K36me3 exhibit distinct downstream effects on gene transcription and chromatin structure (DiFiore et al., 2020). Expression of the NSD2 T1150A variant as abnormal H3K36 trimethyltransferase will lead to increased H3K36me3 levels and introduction of H3K36me3 at aberrant genomic locations.

### 3.8 T1150A mediates upregulation of gene expression via antagonizing H3K27me3

Next, we wanted to get insights into the possible biological effects which could be mediated by NSD2 T1150A in leukemic cancer cells. Bioinformatic analysis of publicly available gene expression datasets was performed to retrieve the change of transcriptome between leukemic cancer cell lines harboring T1150A or WT NSD2. For this, a diffuse large B cell lymphoma (DLBCL) cell line (OCILY-18) was identified in the CCLE database to carry the NSD2 T1150A mutation. OCILY-18 gene expression data were compared with four other DLBCL cell lines containing NSD2 WT revealing a differential gene expression signature with 504 upregulated genes and 163 downregulated genes (Figure 4B, Supplementary Table 2). While these are different cell lines, they are related because all were derived from DLBCL. Control cell lines were selected not to contain mutations in NSD enzymes and SETD2 and not to show overexpression of NSD enzymes in comparison with the other control cell lines. Notably, systematic differential gene expression analyses performed between the four selected NSD2 WT (DLBCL) cancer cell lines (each one against the other three) provided no significant hits (Supplementary Figure 10). To validate that the expression differences of the OCILY-18 cell line from the controls is indeed due to the T1150A mutation, future studies will need to include more T1150A mutant cell lines, or WT and mutant cells with isogenic background.

Assuming that the expression changes of the OCILY-18 cell line are related to the T1150A mutant, we further investigated them to reveal if they contain additional relevant information. The higher number of upregulated genes in the T1150A mutant cell line came in agreement with the known correlation of H3K36me3 and active gene expression. Mapping the differential upregulated and down regulated genes with the COSMIC cancer gene census list (a list of well-established oncogenes and tumor suppressor genes) revealed 15 differentially upregulated oncogenes and 8 downregulated tumor suppressor genes (Figure 4B). For example, HoxA9 was one of the differentially upregulated genes in the tumor cell line containing T1150A, which is a known oncogene in hematological cancers and its expression is under the control of NSD enzymes (Bennett et al., 2017). Due to the lack of H3K36me2/3 ChIP-seq data for these cells, a direct correlation of alterations of these marks and expression changes could not be investigated.

However, there is a strong antagonism between the H3K36me3 and H3K27me3, an important gene silencing modification, which is more prevalent than the antagonism of H3K36me2 and H3K27me3 as illustrated by global mass spectrometric analyses (Leroy et al., 2013; Mao et al., 2015; Voigt et al., 2012). We wondered whether competition of elevated H3K27me3 by the NSD2 T1150A generated H3K36me3 could explain the gene upregulation. To address this question, we analyzed publicly available ENCODE histone modifications ChIP seq datasets for enriched histone modifications at the upregulated genes. Impressively, the H3K27me3 ChIP profile in the B-lymphocyte cell line GM12878 was retrieved as the top significant hit (adjusted p-value 1.32 x 10^-10^) (Figure 4C). Moreover, TF-ChIP seq analysis of upregulated genes revealed an enrichment of EZH2 in B-cell as the only significant hit (adjusted p-value 0.0028) (Supplementary Figure 11A). This finding suggests that these genes are silenced by H3K27me3 in normal B lymphocytes and they become upregulated by the H3K36me3 generated by NSD2 T1150A. H3K27me3 is critical to halt the self-renewal of lymphoid progenitors (Lee et al., 2013; Majewski et al., 2008). Disruption of this function can result in lymphomas similar to the EZH2 inactivating mutants observed in many hematological cancers (Lee et al., 2013; Ntziachristos et al., 2012; Simon et al., 2012) and the results of the H3.3K27M oncohistone frequently observed in pediatric gliomas (Chan et al., 2013; Lewis et al., 2013). Moreover, a decrease in H3K27me3 was correlated with poor outcomes in leukemic patients (van Dijk et al., 2021). Imbalanced high H3K36me3 level can phenocopy the EZH2 disruption by inhibiting its recruitment and activity finally leading to decreased H3K27me3 levels (Yuan et al., 2011).

## 4 Conclusions

In cancer cells, numerous somatic mutations were observed in critical epigenome regulators including the H3K36 dimethyltransferases NSD1 and NSD2. To shed light on possible mechanisms by which these mutations may lead to carcinogenesis, we investigated enzymatic properties of several of somatic missense mutants of NSD2 and NSD1. We show that mutations of Y1971 and R2017 in NSD1 abrogate catalytic activity indicating that NSD1 can act as tumor suppressor gene in some tumor types which is in line with its frequent silencing or deletion in cancers. In contrast, the frequent T1150A mutation of NSD2 and the corresponding T2029A mutant of NSD1 showed hyperactivity and an abnormal H3K36 trimethylation activity in vitro and in human cells, which contrasts the WT NSD1 and NSD2 enzymes that both only function as dimethyltransferases. This novel activity was documented biochemically in vitro and in human cells in a SETD2 knock out background. Using MD simulations, we investigated the mechanistic underpinnings of this atypical activity and uncovered key rules governing the product specificity of this class of enzymes by the accurate positioning of residues surrounding the H3K36 side chain. These analyses revealed an enlarged active site pocket of NSD2 T1150A compared to the WT enzyme in particular in complex with the H3K36me2 substrate, which provides enough space for binding of the AdoMet cofactor allowing the conversion of bound H3K36me2 to the trimethylated product. Additional MD simulations revealed two important contacts of T1150 with Y1092 and L1120 which mediate the change in the active pocket volume between NSD2 WT and the T1150A mutant (Figure 5A). Based on our biochemical, cellular, and computational biology date combined with genome analyses of published data, we propose a model by which this mutant could aberrantly introduce H3K36me3 which competes with endogenous H3K27me3 marks finally leading to changes in gene expression and carcinogenesis (Figure 5B). If this model could be validated in follow up studies in cancer cells, an NSD2 inhibitors would be a therapeutic option for T1150A containing patients.

## Data accessibility

All biochemical data generated or analyzed during this study are included in the published article and its supplementary files. Supplementary Table 3, Movie 1, modelled structures of NSD2 bound to different peptides and cofactors, source data of the results of the MD analysis, MD simulations codes and analysis scripts are provided on DaRUS (https://doi.org/10.18419/darus-3263).

## Author contributions

M.S.K., P.S., S.W. and A.J. devised the study. M.S.K. conducted the biochemical experiments with help by S.W., T.B., A.B. and P.B. P.S. conducted the MD simulations. M.S.K. conducted the bioinformatic work. A.J. supervised the work. J.P. contributed to the supervision of the MD simulations. M.S.K., P.S., S.W. and A.J. prepared the figures. M.S.K., P.S., and A.J. wrote the manuscript draft. All authors were involved in data analysis and interpretation, and preparation of the final manuscript, which was approved by all authors.

## Acknowledgements

This work has been supported by the Deutsche Forschungsgemeinschaft under Germany’s Excellence Strategy EXC 2075 390740016 in PN2-5. M.S.K. was supported by the GERLS scholarship program funding number 57311832 by the German Academic Exchange Service (DAAD) and the Egyptian Ministry of Higher Education.

