## Supplemental Information for "The T1150A cancer mutant of the protein lysine dimethyltransferase NSD2 can introduce H3K36 trimethylation"

### Supporting Information

#### Supplementary Figures

Supplementary Figure 1: Quality control experiments for recombinant H3.1 nucleosome formation
Supplementary Figure 2: Secondary structure analysis of NSD WT and mutant enzymes using CD spectroscopy
Supplementary Figure 3: Additional data regarding H3.1 protein and nucleosome methylation by NSD2 and NSD1 WT and T1150A or T2029A mutants
Supplementary Figure 4: Quality control and validation of H3K36me3 antibody (ab9050)
Supplementary Figure 5: NSD1 methylation assays using the H1.5 (160-176) peptide as substrate
Supplementary Figure 6: Methylation of H3K36me1 peptide by NSD1 T2029A followed by MALDI-TOF mass spectrometry
Supplementary Figure 7: Additional data regarding the MD simulations of NSD2 complexes.
Supplementary Figure 8: Selection and validation of SETD2 KO single HEK293 cell clones
Supplementary Figure 9: Flow cytometry data and analysis of the transfection of different mVenus constructs in SETD2 KO HEK293
Supplementary Figure 10: Differential gene expression analysis of B-cell lymphoma cancer cell lines having NSD2 WT using GEO57083 dataset
Supplementary Figure 11: TFs ChIP-seq analysis for differentially regulated genes in NSD2 T1150A containing cells
Supplementary Figure 12: Criteria used for definition of a successful docking event derived
Supplementary Figure 13: Volume calculation in the active site of NSD2 around K36

#### Supplementary Tables

Supplementary Table 1: List of sgRNA sequences used to target SETD2 in Crispr-Cas9 knockout
Supplementary Table 2: List of peptides used in this study
Supplementary Table 3: List of differentially expressed genes identified in the OCILY18 B-cell lymphoma tumor cell line containing NSD2 T1150A

#### Supplementary Data

- Movie 1: Example of a successful docking of AdoMet to the NSD2 T1150A – H3K36me2 peptide complex.
- Modelled structures of NSD2 bound to different peptides and cofactors
- Source data of the results of the MD analysis
- MD simulations codes and analysis scripts

These data are provided on DaRUS (https://doi.org/10.18419/darus-3263).

### Supplementary Figures

#### Supplementary Figure 1: Quality control experiments for recombinant H3.1 nucleosome formation.

**(A)** Coomassie stained SDS-gel of histone octamers after size exclusion chromatography showing the successful histone octamer formation. **(B)** Electrophoretic mobility gel shift assay (EMSA) of free DNA and nucleosomal DNA showing the successful nucleosome formation.

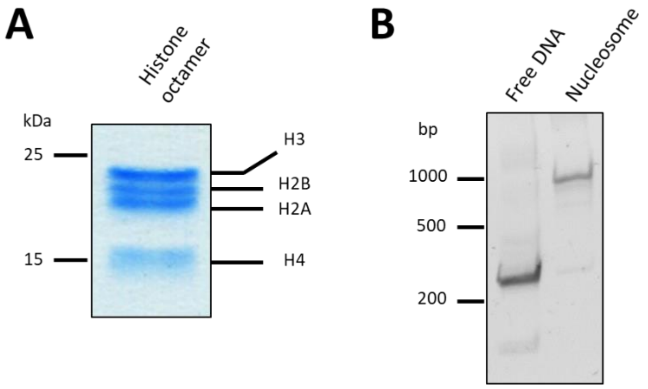

**Supplementary Figure 2: Secondary structure analysis of NSD WT and mutant enzymes using CD spectroscopy.**

**(A)** CD spectra of NSD2 WT compared to the T1150A cancer mutant. **(B)** CD spectra of NSD1 WT compared to the different cancer mutants (Y1971C, R2017Q, R2017L and T2029A). The data document that all mutant proteins are folded. Only in case of the R2017L mutant, slight deviations between WT and mutant CD spectra were observed.

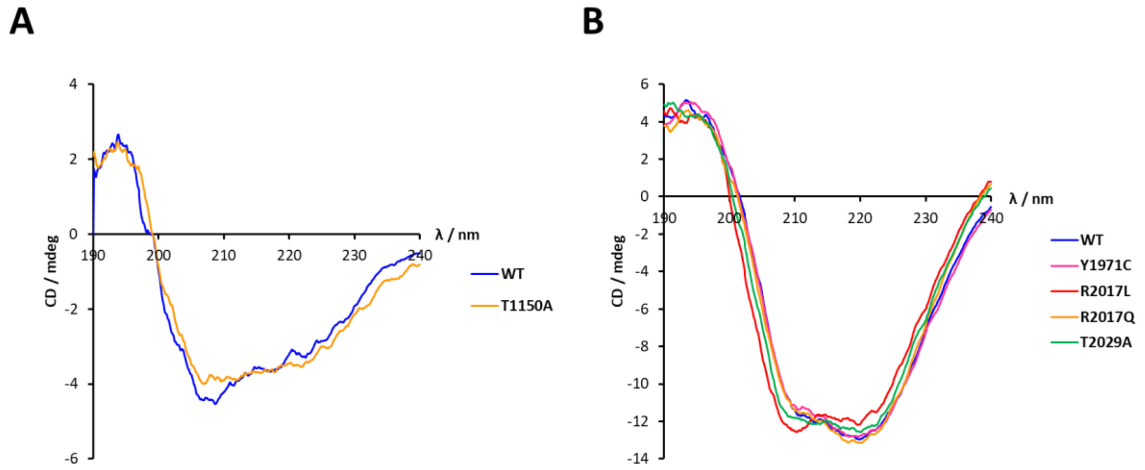

**Supplementary Figure 3: Additional data regarding H3.1 protein and nucleosome methylation by NSD2 and NSD1 WT and T1150A or T2029A mutants.**

**(A)** Western blot analysis of H3K36me2 generation after methylation of H3.1 recombinant nucleosome with NSD2 WT or T1150A. **(B)** Additional example of the H3K36me3 western blot after methylation of H3.1 recombinant protein with NSD1 WT or T2029A (as shown in Figure 2E) with prolonged film exposure. In both panels, the equal loading of substrates and enzymes is shown using Ponceau S staining.

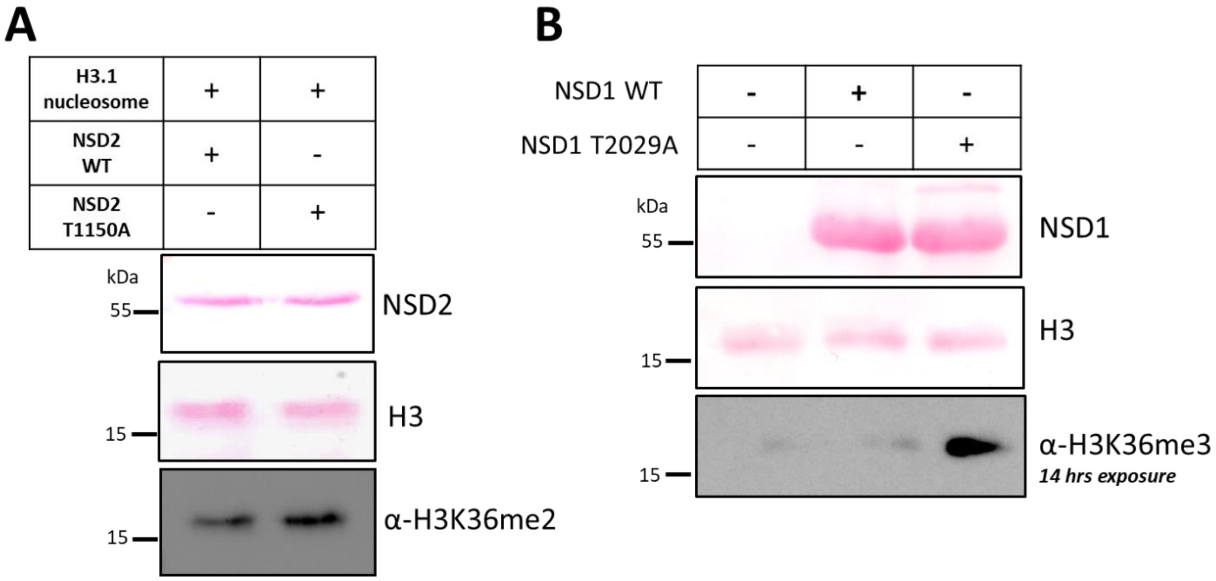

**Supplementary Figure 4: Quality control and validation of H3K36me3 antibody (ab9050).**

**(A)** Binding of the H3K36me3 antibody (ab9050) to a peptide array containing H3 (29-43) peptides with K36me0, me1, me2, me3, and K36A. Binding was detected with HRP-anti rabbit antibody and chemiluminescence signal detection. The antibody shows high specificity for H3K36me3. **(B, C)** Western blot analysis signals after methylation of H3.1 recombinant protein with NSD1 WT or T2029A using the H3K36me2 specific antibody (B) or the H3K36me3 specific antibody (C). The T2029A mutant was used at decreasing concentrations until equal H3K36me2 signals were observed for WT and T2029A as marked by red squares. Under these conditions, only T2029A showed an H3K36me3 signal**.**

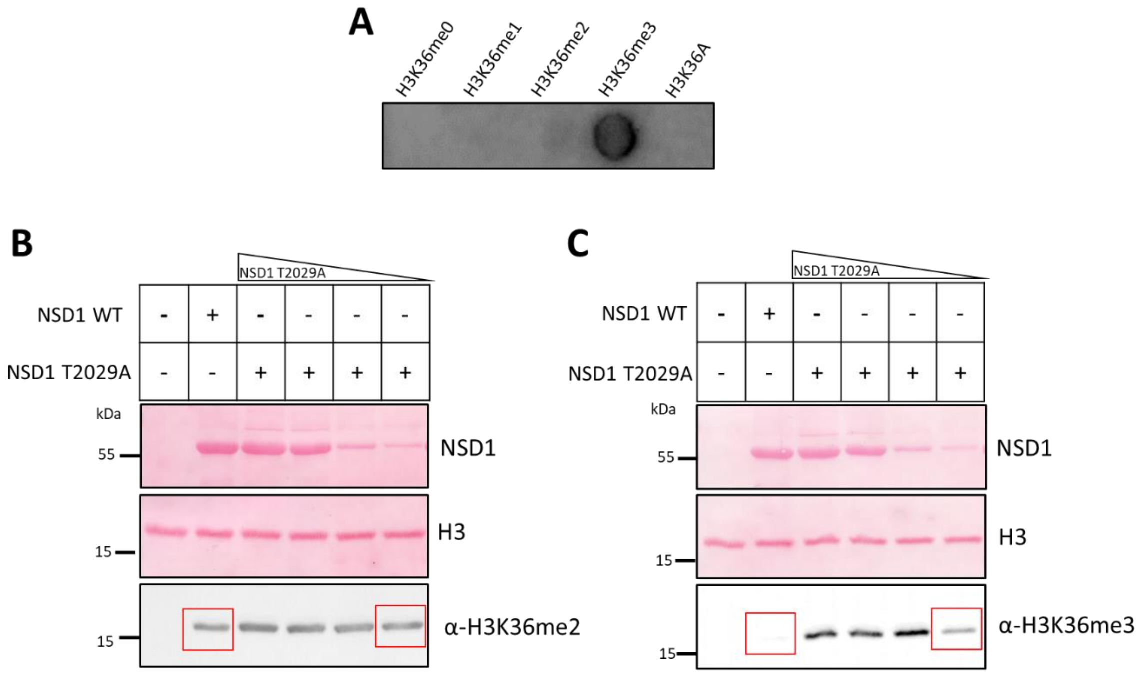

**Supplementary Figure 5: NSD1 methylation assays using the H1.5 (160-176) peptide as substrate.**

**(A)** Methylation of the H1.5 (160-176) peptide containing K168 by NSD1 WT and the different missense cancer mutants using radioactively labelled AdoMet. The upper panel shows the autoradiographic picture and the lower panel depicts the corresponding quantitative analysis showing the mutant activity relative to WT. The data are expressed as means ± SEM for 2 independent replicates. **(B-D)** H1.5 K168 peptide methylation by NSD1 WT or T2029A using unlabeled AdoMet followed by MALDI-TOF mass spectroscopy. H1.5 K168 mass spectra (theoretical mass 2548.38 Da) are shown either without enzyme incubation (B), after incubation with NSD1 WT (C) or T2029A (D). The methylation reaction was incubated for 4 h at 37 °C.

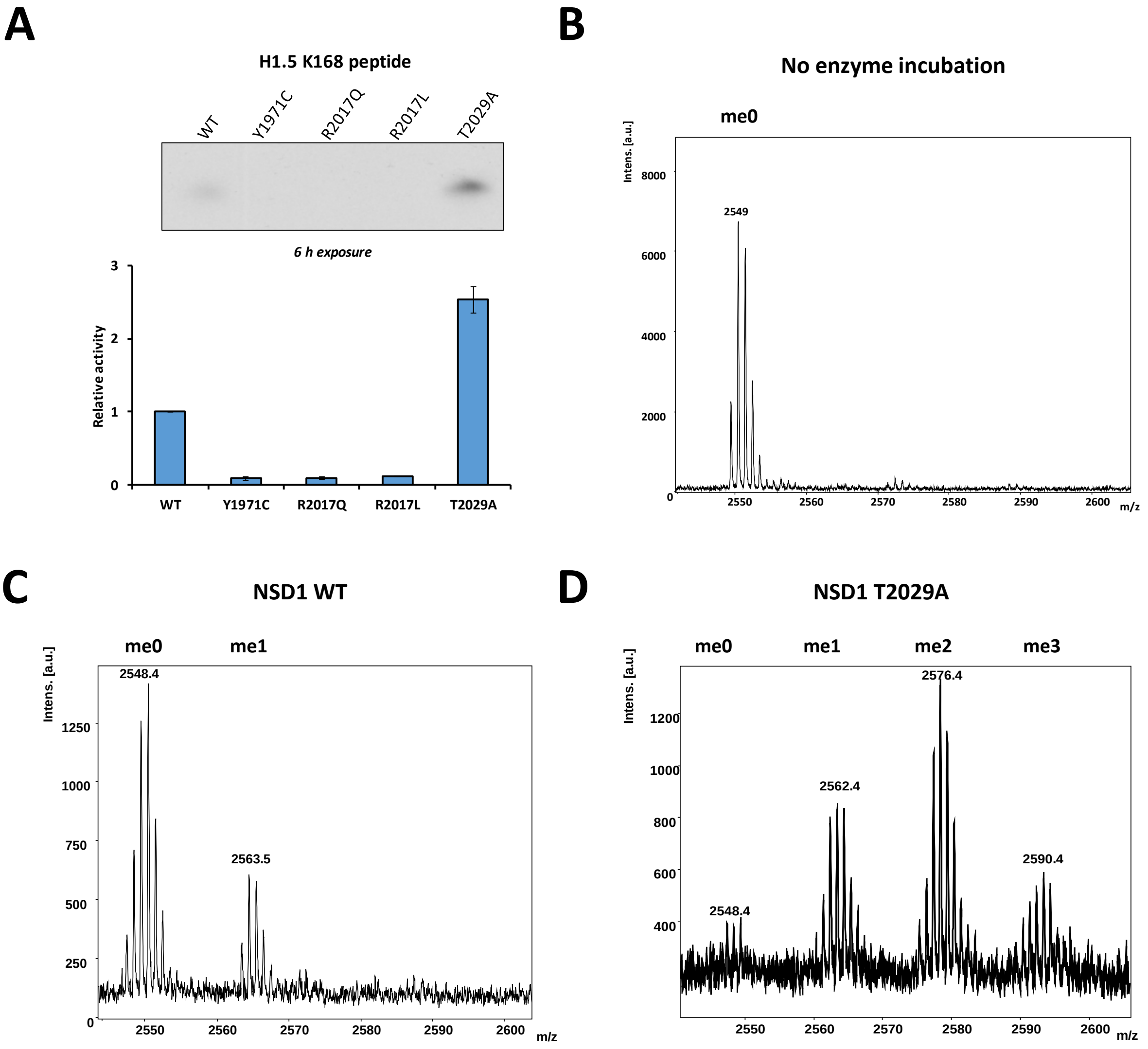

**Supplementary Figure 6: Methylation of H3K36me1 peptide by NSD1 T2029A followed by MALDI-TOF mass spectrometry.**

**(A)** Mass spectrum of the H3K36me1 peptide (theoretical mass MH+ 2118.21) without enzyme incubation. **(B)** Mass spectrum of the H3K36me1 peptide after 4 h incubation with NSD1 T2029A at 37 °C in methylation buffer supplemented with unlabeled AdoMet.

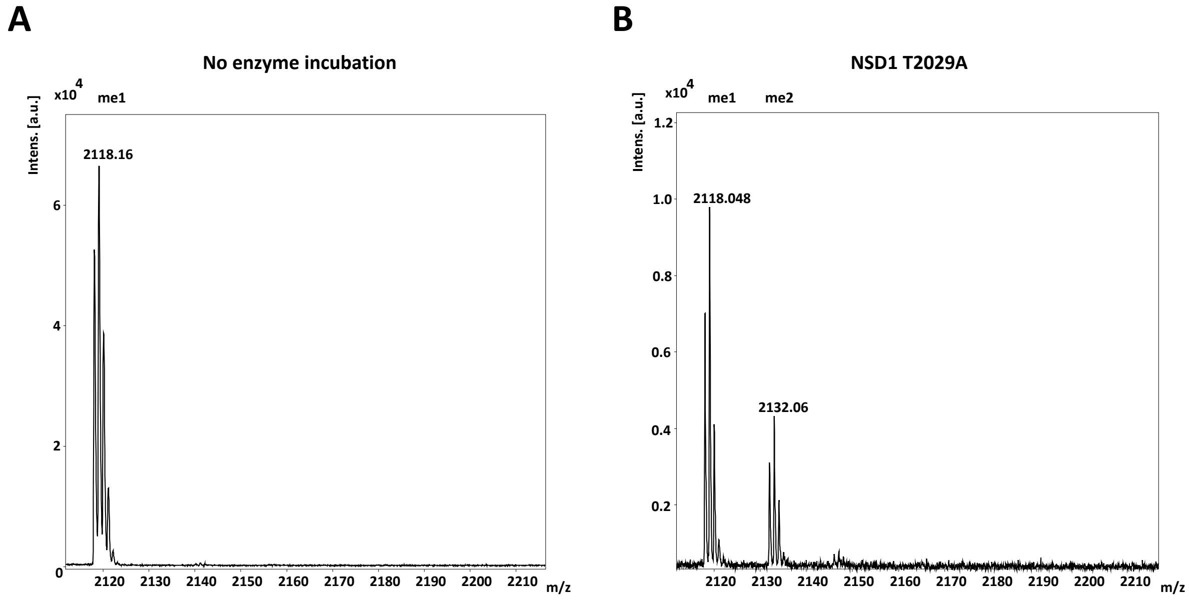

**Supplementary Figure 7: Additional data regarding the MD simulations of NSD2 complexes.**

**(A)** Comparison of the positions and conformations of AdoMet after successful sMD association of AdoMet to NSD2 T1150A – H3K36me2 peptide complexes and AdoMet of the NSD2-peptide-AdoMet complex modelled on the basis of available NSD2-Peptide-AdoHcy structures. The figure shows all atom RMSD values of AdoMet taken from successful AdoMet associations observed in 30 sMD simulations à 100 ns. The RMSD values in the small Å range document similar positions and orientations of AdoMet in both settings. **(B)** Visualization of example structures. The RMSD value in Å is indicated at the top. AdoMet from sMD simulations is shown in purple, AdoMet directly modelled into the complex is shown in orange. **(C)** Enlarged image of the T1150 contact to L1120 and Y1092 taken from one snapshot of MD simulations of NSD2 - H3K36 peptide – AdoMet complexes.

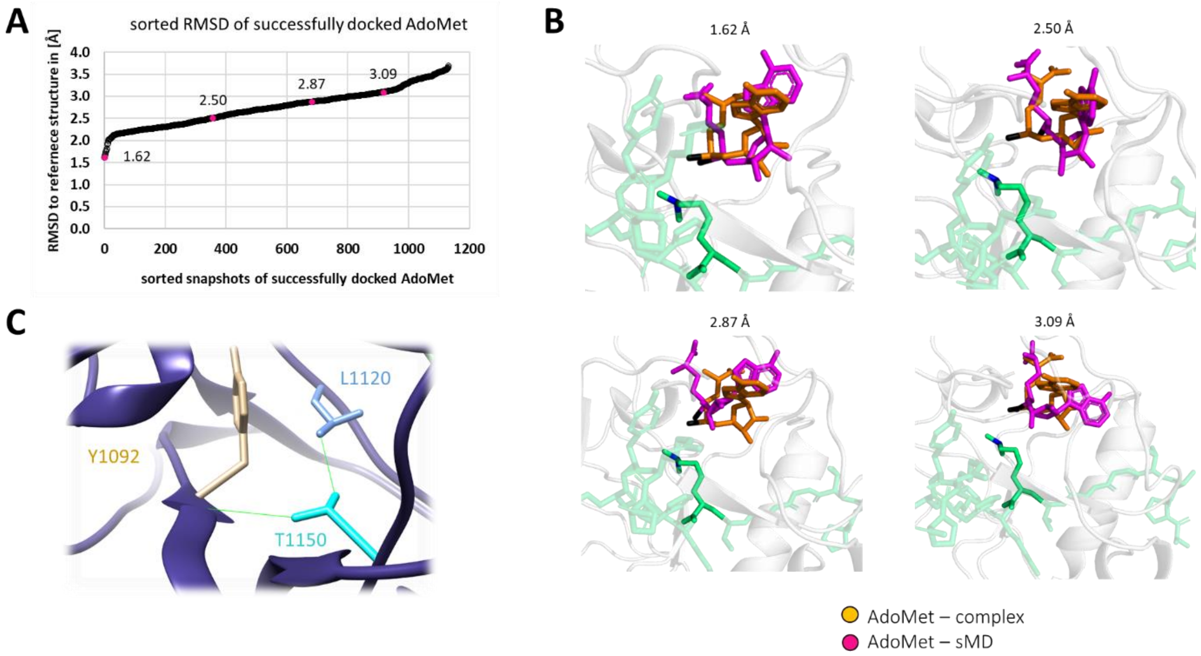

**Supplementary Figure 8: Selection and validation of SETD2 KO single HEK293 cell clones.**

**(A)** Immunoblot analysis of global H3K36me3 contents in cell lysate of HEK293 cells and 4 different SETD2 KO HEK293 single cell clones using the H3K36me3 specific antibody. The data show the complete loss of H3K36me3 in the KO cells. **(B)** 1% agarose gel stained with GelRed showing the PCR products amplified from genomic DNA of SETD2 KO cell clone #4 at exon 3 and 9 of the *SETD2* gene which are targeted by sgRNA #2 and #3, respectively. The figure shows deletions in both alleles of exon 3, insertion at one allele of exon 9 while the second allele of exon 9 showed a similar size as the WT product. **(C)** Sanger sequencing of the PCR product amplified from exon 9 of genomic DNA of SETD2 KO cell clone #4 as shown in panel B in comparison to parental HEK293 cells. The sequence alignment shows the insertion of one guanine base causing a frame shift mutation in this allele of the *SETD2* gene.

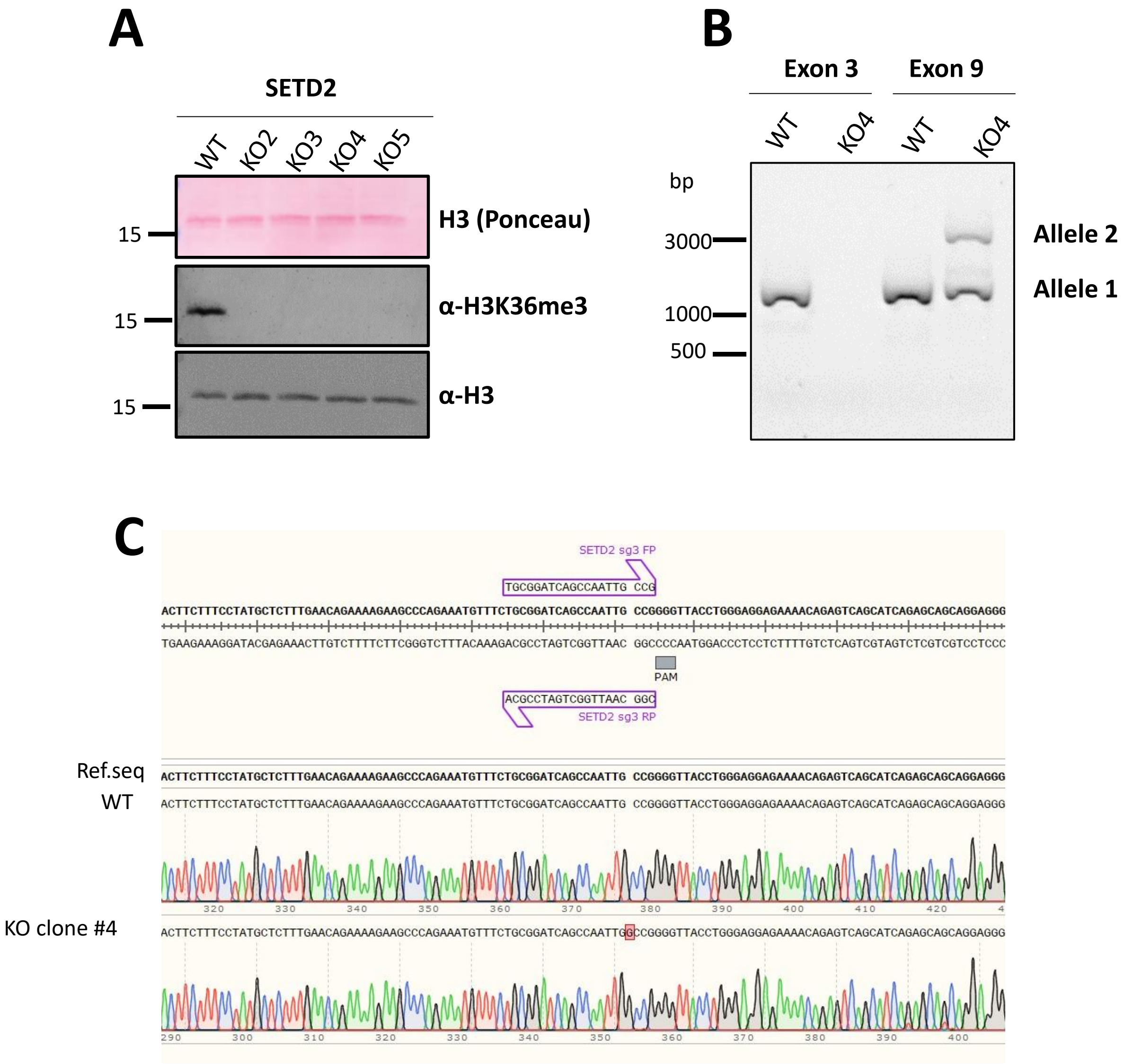

**Supplementary Figure 9: Flow cytometry data and analysis of the transfection of different mVenus constructs in SETD2 KO HEK293.**

**(A)** Transfection of mVenus-tagged NSD2 WT, NSD2 T1150A or mVenus empty vector (EV) into SETD2 KO HEK293 cells. The figure shows an example of the gating strategy applied in the flow cytometry experiments. **(B)** Representative example of the flow cytometry primary data for mVenus expression. **(C)** Average percentage of mVenus positive cells in three independent experiments normalized to mVenus-NSD2 WT. EV, empty vector. Data represent mean ± SEM.

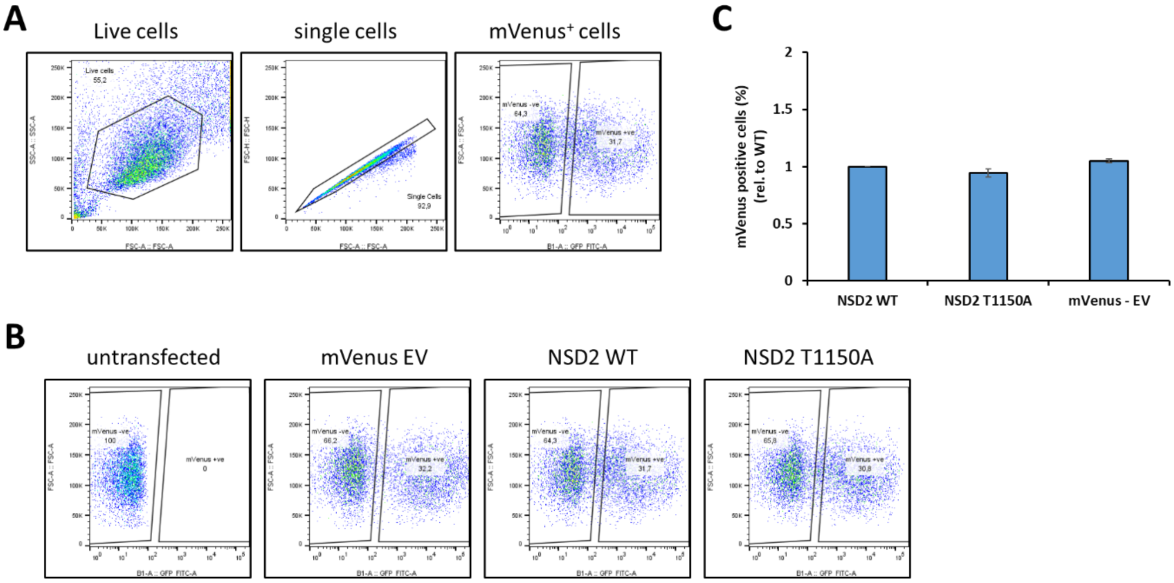

**Supplementary Figure 10: Differential gene expression analysis of B-cell lymphoma cancer cell lines having NSD2 WT using GEO57083 dataset.**

**(A-D)** Volcano plots showing the differential gene expression analysis between 4 B-cell lymphoma cancer cell lines (OCILY-1, OCILY-7, OCILY-10 and DOHH2) all containing WT NSD2 which were used for comparison with OCILY18 containing NSD2 T1150A in Figure 4B. Gene expression of each cell line was compared against the remaining 3 cell lines. Each dot represents one gene probe. There is no significant difference with an adjusted p-value <0.05, hence no green and red dots appear. The analysis was performed by GEO2R tool using GEO57083 expression array data.

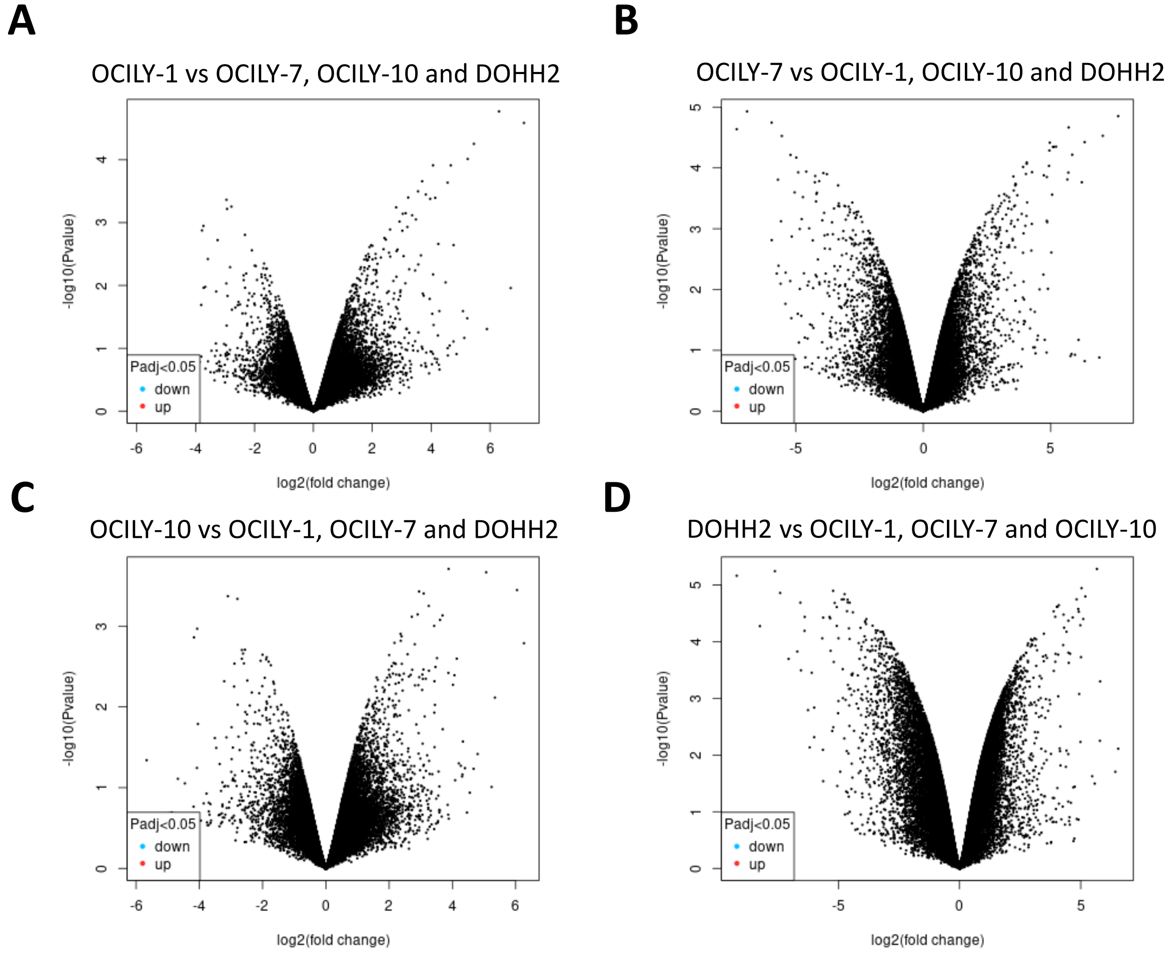

**Supplementary Figure 11: TFs ChIP-seq analysis and gene ontology pathway enrichment analysis for differentially regulated genes in NSD2 T1150A containing cells.**

**(A-B)** TFs ChIP-seq analysis for differentially upregulated (A) and downregulated (B) genes shown in Figure 4B. The figures show the first 3 most significant hits ordered according to their adjusted p-value as indicated at the corresponding bar. Analysis was done using the ENRICHR tool using ENCODE-TF ChIP-seq project data. **(C-D)** GO-biological processes enrichment for differentially upregulated (C) and downregulated (D) genes. The figures show the first 10 most significant hits for upregulated genes (only nine which were significant for downregulated genes). Analysis was done using the ENRICHR tool. All hits are ordered in bar diagram according to their adjusted p-value as indicated at the corresponding bars.

A

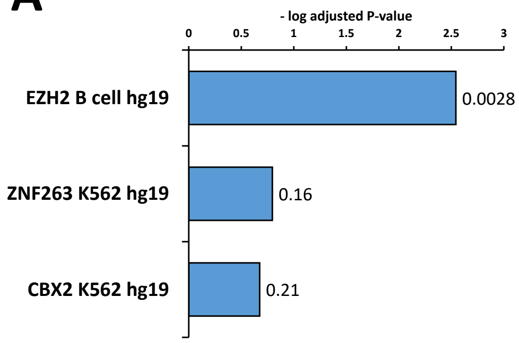


B

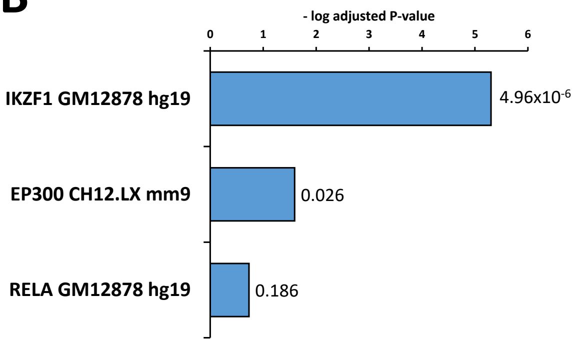

**Supplementary Figure 12: Criteria used for definition of a successful docking event derived from the $S_N2$ TS-like conformation.**

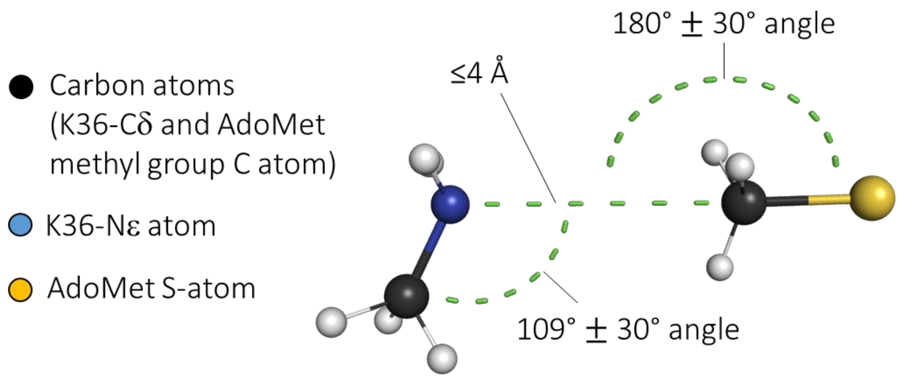

### Supplementary Figure 13: Volume calculation in the active site of NSD2 around K36.

**(A)** Structure of the complex of NSD2 with bound H3K36me2 and AdoMet as well as the inclusion spheres defining the sampled space. **(B)** Raw data of the volume calculations shown in Figure 3D.

**A**

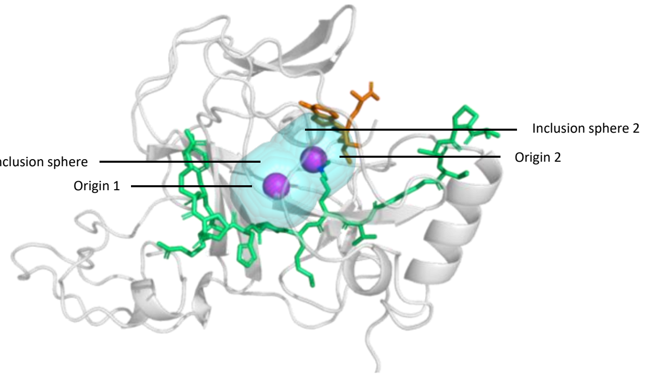


**B**

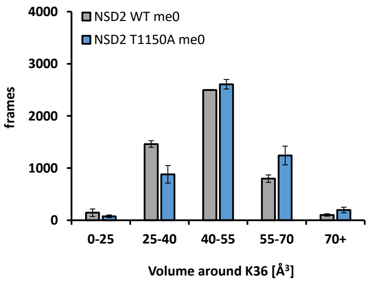


**C**

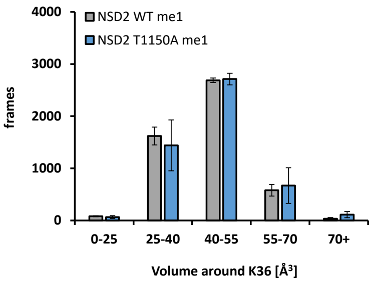


**D**

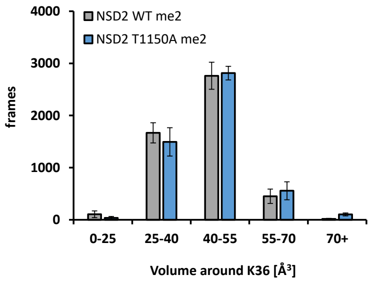

### Supplementary Tables

#### Supplementary Table 1: List of sgRNA sequences used to target SETD2 in Crispr-Cas9 knockout.

| sgRNA number | targeted SETD2 exon | sgRNA sequence |
|---|---|---|
| **1** | 1 | 5′- CCGCAGCCGCCTCCGAAGAT –3′ |
| **2** | 3 | 5′- AATGAACTGGGATTCCGACG –3′ |
| **3** | 9 | 5′- TGCGGATCAGCCAATTGCCG –3′ |

#### Supplementary Table 2: List of peptides used in this study.

Masses refer to monoisotopic mass. Purity was analyzed by HPLC. All peptides were purchased at Intavis AG.

| Name | aa number | sequence | mass (Da) | purity |
|---|---|---|---|---|
| **H3K36** | 26-44 | Biot-R K S A P A T G G V K K P H R Y R P G | 2288.24 | 99.0% |
| **H3K36me1** | 26-44 | Ac R K S A P A T G G V $K^{me1}$ K P H R Y R P G NH2 | 2117.21 | 96.6 % |
| **H1.5 K168** | 160-178 | Biot K K P A A A G V K K V A K S P K K A K(FITC) NH2 | 2548.38 | 98.4 |

#### Supplementary Table 3: List of differentially expressed genes identified in the OCILY18 B-cell lymphoma cell line containing NSD2 T1150A.

Differentially expressed genes were identified by comparison of gene expression data of OCILY18 (containing NSD2 T1150A) control cell line (OCILY1, OCILY7, OCILY10 and DOHH2, all containing NSD2 WT). They are listed with their corresponding $Log_2$ FC and $-log_{10}$ p-values.

Provided as separate file.

**Supplementary Data**

- Movie 1: Example of a successful docking of AdoMet to the NSD2 T1150A – H3K36me2 peptide complex.
- Modelled structures of NSD2 bound to different peptides and cofactors
- Source data of the results of the MD analysis
- MD simulations codes and analysis scripts

These data are provided on DaRUS (https://doi.org/10.18419/darus-3263).
